# MICROGLIAL EXTRACELLULAR VESICLES MEDIATE C1Q DEPOSITION AT THE PRE-SYNAPSE AND PROMOTE SYNAPTIC PRUNING

**DOI:** 10.1101/2025.02.28.640756

**Authors:** Giulia D’Arrigo, Giulia Cutugno, Maria Teresa Golia, Francesca Sironi, Marta Lombardi, Sara Francesca Colombo, Roberto Frigerio, Marina Cretich, Paola Gagni, Elisabetta Battocchio, Cristiana Barone, Emanuele Azzoni, Sonia Bellini, Claudia Saraceno, Roberta Ghidoni, Caterina Bendotti, Rosa Chiara Paolicelli, Martina Gabrielli, Claudia Verderio

## Abstract

C1q is released by microglia, localizes on weak synapses and acts as a tag for microglial synaptic pruning. However, how C1q tags synapses during the pruning period remains to be fully elucidated. Here, we report that C1q is delivered by microglia to pre-synaptic sites that externalize phosphatidylserine through extracellular vesicles. Using approaches to increase or reduce vesicles production in microglia, by C9orf72 knock out or pharmacological inhibition respectively, we provided mechanistic evidence linking extracellular vesicle release to pre-synaptic remodelling in neuron-microglia cultures. In C9orf72 knockout mice, we confirmed larger production of microglial extracellular vesicles, and showed augmented C1q presynaptic deposition associated with enhanced engulfment by microglia in the early postnatal hippocampus. Finally, we provide evidence that microglia physiologically release more vesicles during the period of postnatal circuit refinement. These findings implicate abnormal release of microglial extracellular vesicles in both neurodevelopmental and age-related disorders characterized by dysregulated microglia-mediated synaptic pruning.

## INTRODUCTION

Synaptic pruning, i.e. elimination of synapses formed in excess, is a fundamental process for neural circuits refinement during brain development and microglia recently emerged as important players in this process. During synaptic pruning less active or “weak” synapses are removed while appropriate connections are strengthened (Presumey et al., 2017; Schafer et al., 2012). Increased or abnormal synaptic pruning occurs throughout life in neurological diseases and constitutes a critical step in the development of age-related neurodegenerative diseases, characterized by pathological loss of synapses and cognitive decline (Hong et al., 2016; Thion and Garel, 2018). Several microglial receptors contribute to microglia-mediated synaptic engulfment through the recognition of specific signals tagging weak synapses. The fractalkine receptor CX3CR1 (Paolicelli et al., 2011) detects the chemokine CX3CL1, a “find me” signal released by neurons. The complement receptor 3 (CR3), also known as Mac1 (Schafer et al., 2012), detects cleaved complement 3 (C3), an immune molecule which binds to synapses to be removed downstream C1q, the initiating molecule of the classical complement pathway. C1q and C3 serve as “eat me” signals, which promote phagocytosis of unwanted synapses, and are primarily secreted by microglia and astrocytes in the brain (Fonseca et al., 2017; Zhang et al., 2023). The microglial receptor Trem2 (Filipello et al., 2018) and GPR56 (Peet et al., 2020) target synapses which externalize phosphatidylserine (PS), a phospholipid normally facing the cytoplasm, recently identified as a neuronal clue for microglial synapse elimination during development (Li et al., 2021; Peet et al., 2020; Scott-Hewitt et al., 2020). Externalized PS is also bound by C1q, through its globular head (Paidassi et al., 2008), thus acting as an upstream mediator of both complement- and TREM2/GPR56-dependent synapse elimination. Conversely, the microglial receptor SIRPα detects CD47, a surface immune molecule that acts as “don’t eat me signal”, protecting synapses from excessive removal (Lehrman et al., 2018).

Extracellular vesicles (EVs) are nano- to micro-sized membrane vesicles, generated in the endocytic compartment (exosomes) or at the plasma membrane (ectosomes), which are released by all cells. EVs carry a variety of donor cell components in their lumen and at their surface and transport these molecules between cells, playing a key role in intercellular communication (Gassama and Favereaux, 2021).

EVs produced by microglia, especially large EVs shed from the plasma membrane, externalize PS (Bianco et al., 2009; Perez et al., 2023) and carry C1q and other complement factors (Drago et al., 2017; Lombardi et al., 2019), which form a protein corona around EVs with additional surface molecules (Buzas et al., 2018; Toth et al., 2021). Moreover, large microglial EVs move at the neuron surface, more efficiently along axons than dendrites (Gabrielli et al., 2022), to reach yet unidentified EV-neuron contact sites, where microglial EVs make more stable interactions and may deliver their cargoes. Collectively, these findings suggest that microglial EVs may act as delivery vehicles for complement factors to synapses in need to be removed, a still unexplored hypothesis.

By manipulating microglial EVs concentration in neuron-microglia co-cultures through EVs supplementation, the use of mutant microglia (C9orf72 knock out) producing more EVs and a pharmacological inhibitor of EVs release (GW4869), here we show that microglial EVs deliver C1q to pre-synaptic sites and promote microglia-mediated synaptic pruning *in vitro* and *in vivo* in the CA1 hippocampal region during the critical period of postnatal circuit refinement. Furthermore, we show that microglial EVs production temporally correlates with the pruning period during postnatal brain development.

## RESULTS

### Microglial EVs make persistent interaction with synaptic sites and tag synapses with C1q

As a first step to investigate whether microglial EVs might act as delivery vehicles for complement factors to synapses, we first re-assessed the presence of C1q in EVs released by primary murine microglia. We stimulated microglial cells with ATP, to increase EVs production, and isolated EVs by differential centrifugation at *100000x g* after pre-clearing of cell supernatant from cells and debris at *300x g*, as previously described (Lombardi et al., 2019). Western blot (WB) analysis showed that microglial EVs were enriched in the EVs markers Alix, Flotillin I and Annexin-A2 and unstained for EVs negative makers TOM20 and GS28, respectively mitochondrial and Golgi proteins, excluding contamination with intracellular organelles (Fig. 1A and S1 for ponceau). The analysis confirmed that EVs stained positive for C1q, which was enriched in EVs compared to primary microglia (Fig. 1A), in line with previous evidence (Drago et al., 2017; Lombardi et al., 2019). Further C1q quantification by ELISA in EVs and EVs-depleted supernatant showed that EVs contain about 60% of the secreted factor (C1q content: 12.8 pg in EVs; 8.7 pg in EVs-depleted supernatant conditioned by 1X10^6^ cells). According to MISEV guidelines (Welsh et al., 2024) microglial EVs were further characterized by cryo-electron microscopy (cryo-EM) and tunable resistive pulse sensing (TRPS), that measures EV size and concentration. These analyses showed that most EVs were round in shape and free of contaminants (protein aggregates/intracellular organelles) (Fig. 1B), and had a mean size of 186.8 ± 11nm and mode of 130.6± 7.8nm (Fig. 1C).

**Figure 1.**
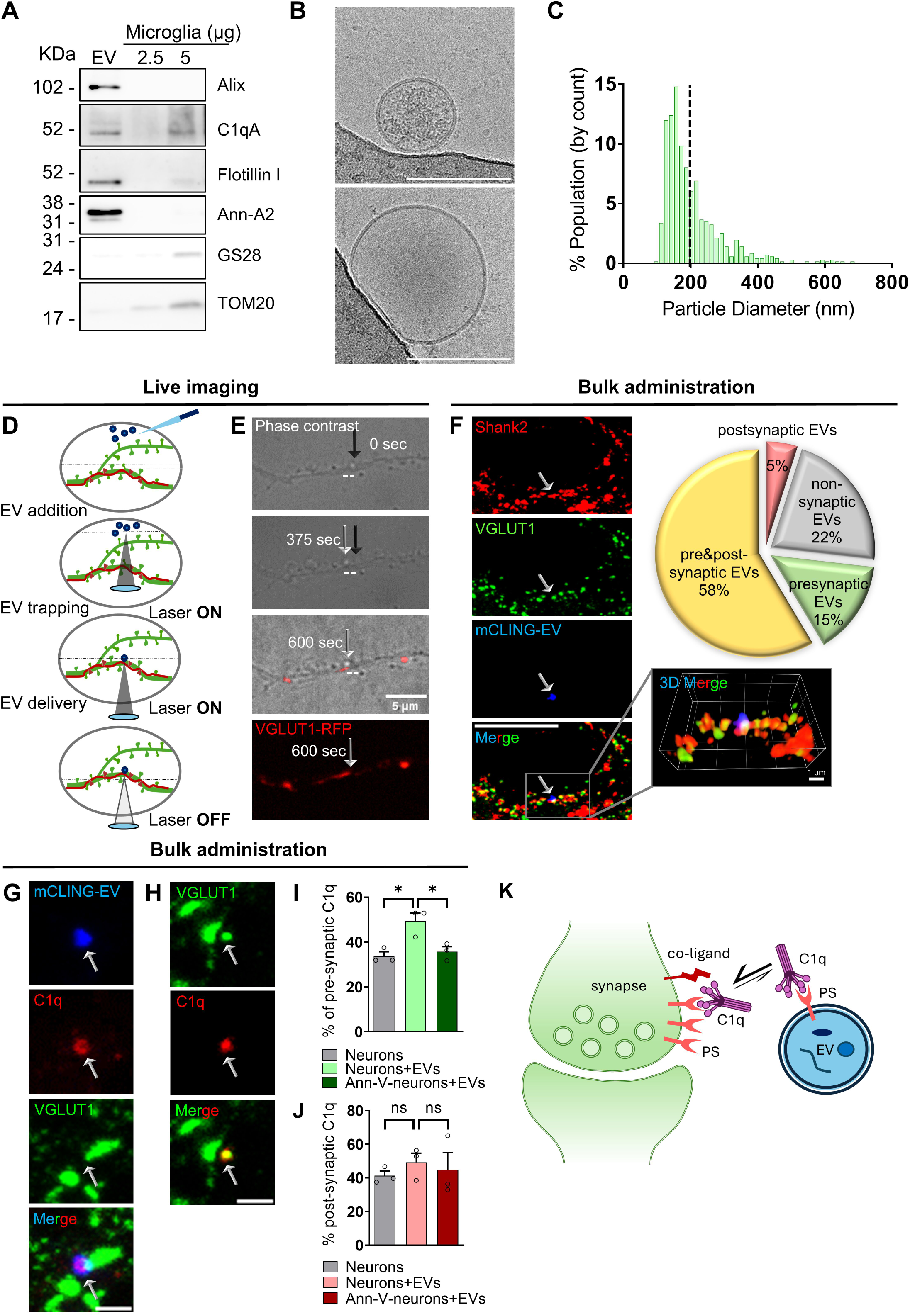
Large microglial EVs localize at synaptic sites and tag synapses with C1q. **A**) Western blotting analysis of microglial EVs in the *100000x g* pellet from 1 × 10^7^ microglia and donor cell lysate (2.5-5µg) for the EVs markers Alix and Annexin-A2 (Ann-A2), Flotillin I, the mitochondrial marker TOM20, the Golgi protein GS28 and the complement factors C1q. Ponceau staining is shown in Fig. S1. **B**) Representative cryo-EM micrographs of microglial EVs in the *100000x g* pellet. Scale bar 100 nm **C**) Representative size distribution of EVs analysed by TRPS (number of independent experiments (N)=3). **D**) Schematic representation of large EVs (≥ 200 nm) delivery to dendrites of mature hippocampal neurons using optical tweezers. A single EV was trapped by the laser tweezers in bright field and placed in contact with the dendrite receiving VGLUT1-RFP puncta. The trapping laser was switched off 30 seconds after contact and EV-dendrite interaction was monitored in bright field for 10 minutes. **E**) Sequence of phase contrast, confocal fluorescence or merged images at 0, 375 and 600 seconds after contact. Black arrow indicates the EV position at 0 sec, grey arrow the EV position at 375 and 600 sec. Dotted line indicates EV displacement. Scale bar 5 μm. Number of EVs which stopped at VGLUT1+ puncta n=7/14. **F**) Representative confocal acquisition of a mCLING-labelled EV (blue) interacting with VGLUT1+ (green) and Shank2+ (red) puncta at 3 hours after addition to neurons. Scale bar 10 μm. Zoom-in: 3D reconstruction by Imaris, scale bar 1 μm. Right, pie chart showing the percentage of mCLING-labelled EVs with pre- and/or post-synaptic localization (n=41 analysed EVs). **G**) Representative confocal image of a mCLING-labelled EV (blue) positive for C1q (red) interacting with a VGLUT1+ (green) terminal. Scale bar 5 μm. **H**) Representative confocal acquisition of a C1q spot (red) co-labelled by VGLUT1 (green) in neurons exposed to EVs. Scale bar 5 μm. **I**) Percentage of C1q puncta interacting with pre-synaptic terminals in EVs-treated neurons and neurons treated with EVs after Annexin-V (Ann-V) exposure. Each data point represents the percentage of pre-synaptic C1q over total C1q puncta per experiment: number of analyzed C1q puncta=700 from N=3. Holm-Sidak’s multiple comparisons test: Neurons+EVs vs. Neurons p=0.0175; Neurons+EVs vs. Ann-V-neurons+EVs p=0.0214. **J**) Percentage of C1q puncta interacting with PSD95+ terminals as in **I**: number of C1q puncta=787 from N=3. Holm-Sidak’s multiple comparisons test: not significant (ns). K) Scheme depicting the mechanism possibly mediating C1q transfer from EVs to synapses externalizing PS.

To investigate whether EVs might establish persistent interactions with synapses we transfected mouse hippocampal neurons with eGFP and VGLUT1-RFP to delineate dendrites receiving VGLUT1+ puncta at 13 days in vitro (13 DIV), placed single large EVs (above 200 nm in size, Fig. 1C) on dendrites receiving VGLUT1+ puncta using optical manipulation, and monitored EVs dynamics by time lapse imaging (Fig. 1D), as previously established in our laboratory (D’Arrigo et al., 2021; Gabrielli et al., 2022; Prada et al., 2018). Briefly, EVs were added to the neuronal medium by a pipette, a single EV was trapped by optical tweezers and kept in contact with the selected dendrite for 30 seconds. Then the trapping laser was switched off and time-lapse images were collected (Fig. 1D). We found examples of moving EVs which stopped closed to VGLUT1+ terminals (Fig. 1E; N=7/14).

To further explore the capacity of microglial EVs to make stable interactions with synaptic sites, we added fluorescent EVs, labelled with the fluorescent dye mCLING, in bulk to neurons for 3 hours, to allow EVs to reach their target sites on neurons, and then fixed and stained neurons with the pre- and post-synaptic markers VGLUT1 and Shank2 and the complement factor C1q. Immunofluorescence analysis revealed that 78% of mCLING-labelled EVs localized closed to VGLUT1+ and/or Shank2+ terminals (Fig. 1F, number of analysed EVs n=41), with no preference for pre- or post-synapses (Fisher’s test p=0.2565), and that about one-third of the EVs were C1q+ (Fig. 1G, number of C1q+EVs=42/145). Quantification of C1q puncta adjacent to VGLUT1+ terminals (Fig. 1G) and/or co-labelled with VGLUT1 (Fig. 1H) revealed a higher percentage of C1q-tagged pre-synapses (+47%) in neurons exposed to EVs compared to untreated neurons (Fig. 1I). About half of pre-synapses tagged by EVs-derived C1q were mCLING negative (Fig. S2), suggesting that microglial EVs can deliver surface C1q to pre-synaptic sites. Pre-treatment of neurons with annexin-V to cloak externalized PS, a known C1q ligand (Paidassi et al., 2008), prevented EVs-mediated C1q deposition to pre-synaptic sites (Fig. 1I). By contrast, we found no significant changes in the percentage of C1q puncta interacting with PSD95+ post-synaptic terminals in EVs-treated neurons (Fig. 1J). These findings indicated that microglial EVs deliver surface C1q to pre-synapses that externalize PS. For this to occur, C1q should detach from PS exposed on the EV and bind to PS externalized on the pre-synapse, a process that might be favoured by larger amount of PS and/or the presence of C1q co-ligand(s) on the pre-synapse (Fig. 1K).

### Microglial EVs promote synaptic pruning

Having found that microglial EVs interact with neurons at synapses and tag them with C1q, we next asked whether EVs could promote synaptic pruning. We supplemented EVs in bulk to neurons, then plated microglia on the culture (at 1-to-1 microglia-to-neuron ratio) and kept the cells in co-culture for 24 hours. Finally, we fixed and stained the cells with the microglial marker IB4 and the pre- and post-synaptic markers Bassoon and Shank2, respectively, to quantify synaptic density and microglial synaptic engulfment (Fig. 2A). Neurons cultured alone, exposed or not to EVs, and neurons cocultured with microglia not exposed to EVs were used as controls. Confocal acquisition and analysis of Bassoon+ and Shank2+ puncta along dendrites close to microglia (from 10 to 50 μm) revealed normal pre-synaptic density (Fig. 2B-C) but lower post-synaptic density (Fig. 2B, D) in neurons co-cultured with microglia compared to neurons alone. A decrease in pre-synaptic density was instead observed in EVs-treated neurons co-cultured with microglia (Fig. 2C), with no further decrease in post-synaptic density (Fig. 2D), suggesting that EVs may drive pre-synaptic microglial removal. No changes in synaptic density were induced by EVs on neurons cultured alone (EVs-treated neurons normalized on neurons alone: mean pre-synaptic density=1.035, Mann-Whitney test: p=0.5880, mean post-synaptic density=1.008, Mann-Whitney test: p=0.4463, number of fields >50/condition, N=4). Consistent with these findings, confocal acquisition and imaging analyses revealed that IB4+ microglia displayed higher engulfment of Bassoon+ pre-synaptic puncta when cultured with neurons pre-exposed to EVs compared to untreated neurons (Fig. 2E). To corroborate the capacity of microglial EVs to induce synaptic pruning, we next isolated synaptosomes from the mouse brain, incubated or not synaptosomes with microglial EVs for 3 hours, added synaptosomes to microglia and finally fixed and stained the cells with VGLUT-1 for quantification of their uptake (Fig. 2F). The analysis revealed that microglia engulfed more EVs-treated synaptosomes compared to untreated ones (Fig. 2F), confirming that EVs facilitate engulfment of synaptic terminals. Quantification of Shank2+ material revealed more Shank2+ postsynaptic puncta inside microglia cultured with neurons pre-exposed to EVs compared to untreated neurons (Fig. 2E), suggesting that pre-synaptic engulfment is accompanied by postsynaptic removal.

**Figure 2.**
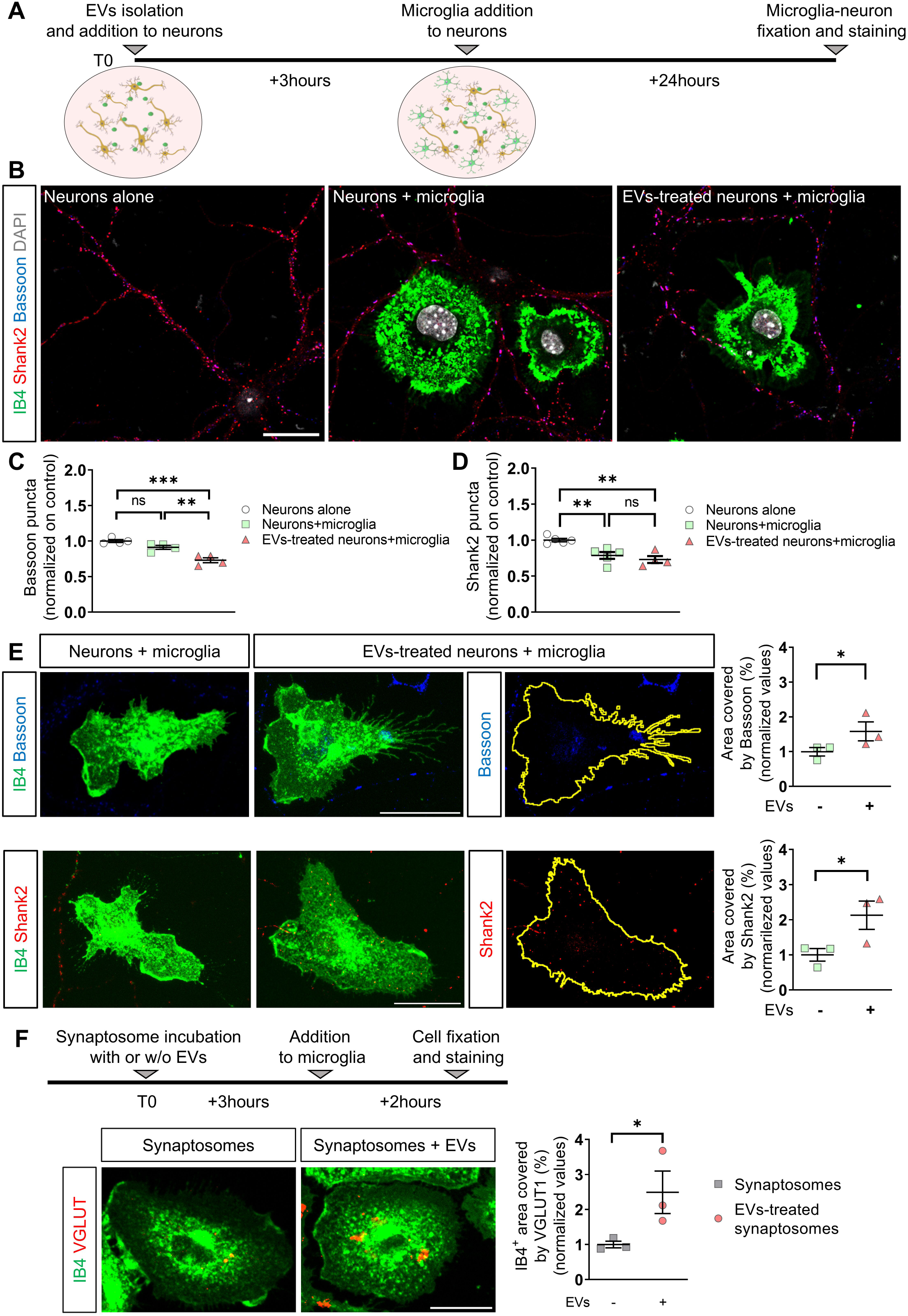
EVs supplementation to neurons drives pre-synaptic pruning. **A**) Schematic of experimental design. **B**) Representative confocal acquisition of neurons alone and neuron-microglia culture pre-exposed or not to EVs (scale bar 20 μm), stained by IB4 (green) and for Shank-2 (red), Bassoon (blue) and nuclei (grey). **C-D**) Relative quantification of pre- and post-synaptic density. Each data point represents mean synaptic density normalized on neurons alone per independent experiment: n=150 analyzed dendrites/condition from N≥4. Holm-Sidak’s multiple comparisons test, (pre-synaptic): Neurons alone vs. EVs-treated neurons+microglia p=0.0003, Neurons+microglia vs. EVs-treated neurons+microglia p=0.0029; (postsynaptic): Neurons alone vs. Neurons+microglia p=0.0045, Neurons alone vs. EVs-treated neurons+microglia p=0.0019. **E**) Representative z-stack projections of IB4+ microglia cocultured with control neurons or EVs-treated neurons (left and middle images) and corresponding quantification of IB4+ area covered by Bassoon or Shank2 (yellow line, right images). Scale bar 20 μm. Data point represent the mean percentage of Bassoon+/Shank2+ area over IB4+ area per experiment normalized on untreated neurons: n=45 analyzed fields per condition from N=3. Basson, Mann-Whitney test, p=0.05. Shank2: Mann-Whitney test, p=0.05. **F**) Top, schematic of experimental timeline. Left, representative z-stack projections of IB4+ microglia (green) that has engulfed vGLUT1+ synaptosomes (red), pre-treated or not with EVs (scale bar 20 μm). Right, relative quantification of the microglial area covered by VGLUT1. Data points represent the mean percentage of VGLUT1+ over IB4+ microglial area normalized on untreated synaptosomes per experiment: n=45 analyzed fields per condition from N=3, Mann-Whitney test p=0.036.

### C9orf72 knock out postnatal microglia release more EVs and complement factors while displaying normal phagocytic activity

To assess the capacity of EVs to enhance microglial synaptic engulfment in a more physiological setting, avoiding EVs supplementation to neurons, we then looked for mutant microglia producing more EVs but displaying normal phagocytic activity, to be co-cultured with neurons for analysis of synaptic pruning. Extensive literature indicates that lysosomes accumulation is often associated with enhanced production of EVs (Eitan et al., 2016), a mechanism for the cells to get rid of unwanted material. Thus, we focused on microglia lacking chromosome 9 open reading frame 72 (C9orf72), a lysosomal regulator (Bauer et al., 2022; Beckers et al., 2021), the absence of which induces accumulation of lysosomes in adult mouse microglia (Lall et al., 2021; O’Rourke et al., 2016). In humans, reduced C9orf72 expression is caused by hexanucleotide repeat expansions in the C9orf72 gene, the most common genetic causes of familial frontotemporal lobar degeneration (FTLD) and amyotrophic lateral sclerosis (ALS) (DeJesus-Hernandez et al., 2011; Renton et al., 2011).

We first investigated whether C9orf72 deficiency may cause lysosomal accumulation in postnatal microglia *in vitro*. Immunofluorescence and WB analysis indicated higher expression of lysosomal proteins (Lamp-1, Cathepsin D, CD68) in C9orf72 knock out (KO) microglia compared to wild type (WT) cells (Fig. 3A-C). We then quantified EVs production using TRPS and found that C9orf72 KO microglia release twice the amount of EVs compared to WT cells (Fig. 3D). EVs production was rated to 3.64x10^7^±9.35x10^6^ per million cells in 1 hour in C9orf72 KO microglia (N=4), and to 1.69x10^7^±4.69x10^7^ EVs per million cells in 1 hour in WT microglia (N=4) (Fig. 3D). C9orf72 KO microglia released EVs larger in size compared to WT cells (Fig. 3D; Mann-Whitney test p=0.036), suggesting increased production of quite large EVs budding from the plasma membrane. Consistent with higher EVs production, KO microglia released more C1q in association with EVs, as evidenced by WB analysis of EVs produced by an equal number of KO and WT microglia (Fig. 3E). When normalized to the EVs marker Alix, the content of C1q was similar in mutant and WT EVs, indicating that higher complement levels in KO EVs reflected larger EVs production rather than complements sorting (Fig. 3E). Importantly, lack of C9orf72 did not alter surface microglial expression of CD11b, the α chain of C3R, the phagocytic receptor mediating complement dependent synaptic pruning, as evidenced by immunofluorescence analysis of mutant and WT cells (Fig. 3F). Of note, C9orf72 did not alter the phagocytic capacity of microglia, as evidenced by analysis of engulfment of fluorescent beads (Fig. 3G) or pHrodo-labelled synaptosomes at 4 hours after incubation (T0, Fig. 3H). Conversely, C9orf72 KO microglia exhibited enhanced degradative capacity, as evidenced by quantification of synaptosome degradation at 4 hours after washout: a lower amount of engulfed synaptosomes was detected in C9orf72 KO microglia compared to WT cells (T4, Fig. 3H). Collectively, these findings identified postnatal C9orf72 KO microglia as a useful tool to explore the role of endogenously released EVs in microglia-mediated synaptic pruning.

**Figure 3.**
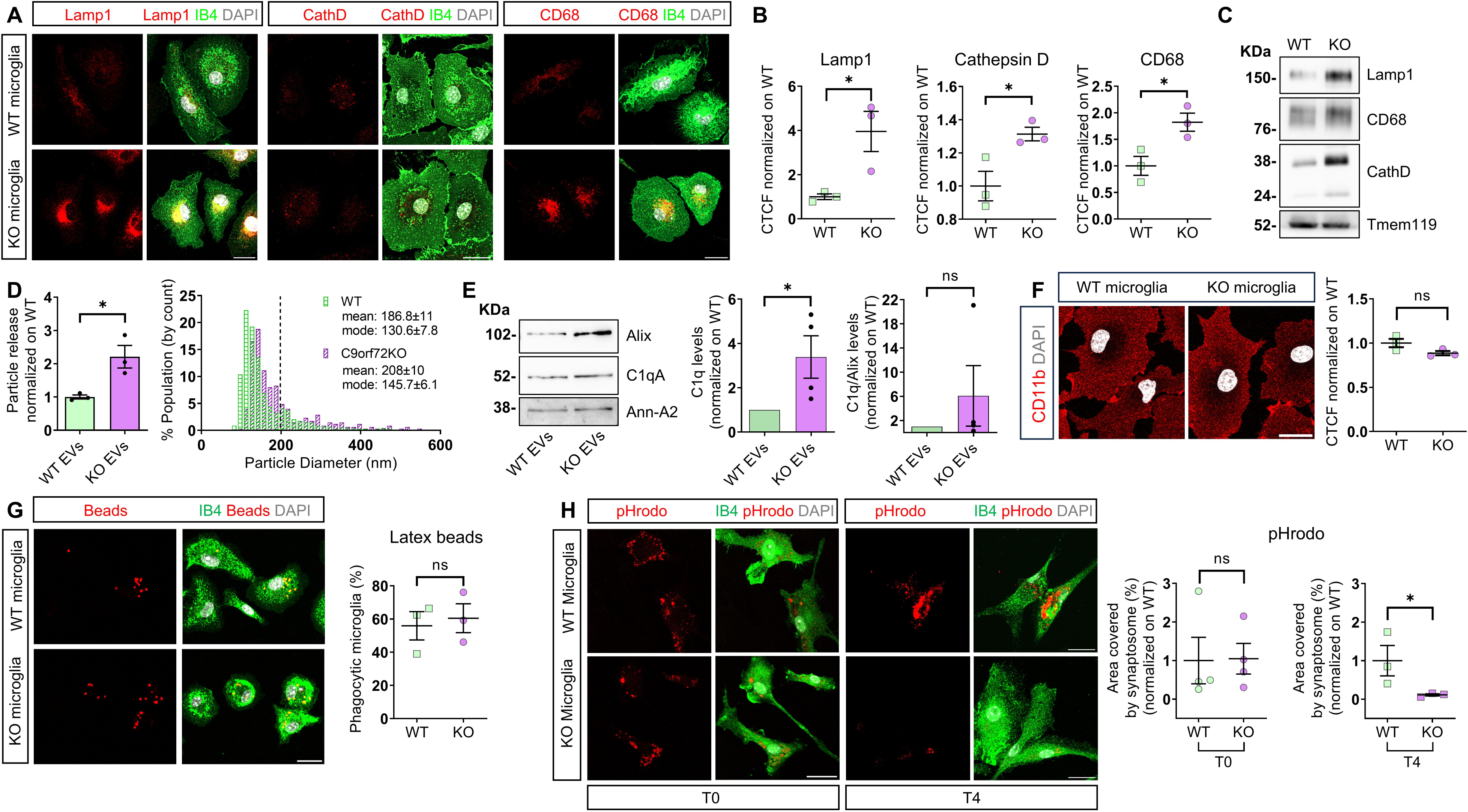
EVs production and phagocytosis by C9orf72 KO microglia. **A**) Representative confocal images of WT and C9orf72 KO microglia identified by IB4 (green) and stained for the lysosomal protein Lamp1, Cathepsin D (CathD) or CD68 (red) and nuclei (grey). Scale bar 20 µm. **B**) Relative quantification of lysosomal proteins. Each data point represents mean corrected total cell fluorescence (CTCF) normalized on WT per experiment : n of analyzed cells ≥35/genotype from N=3; Unpaired t test: Lamp1 p=0.0323, CathD p=0.0330, CD68 p=0.0288. **C**) Representative WB analysis of WT and KO microglia for lysosomal markers and Tmem119. **D**) Constitutive EVs production from 1X10^6^ WT and C9orf72 KO microglia analysed by TRPS (left). Each data point represents single measurement from N=3, normalized on WT cells. Unpaired t test p=0.0247. Size distribution of EVs from WT and C9orf72 KO microglia (right). **E**) Representative WB for C1q of EVs released from 20×10^6^ WT and C9orf72 KO microglia (left) and quantification of vesicular C1q normalized on WT EVs, N=4 (middle, Unpaired t test: p=0.0470) or Alix (right, Mann Whitney test: p=0.3143). **F**) Representative confocal images of WT and C9orf72 KO microglia stained for CD11b (scale bar 20 µm) and relative quantification of CTCF. Each data points represent mean CTCF normalized on WT per experiment: n of analyzed cells=60 /genotype from N=3. Unpaired t test p=0.0992. **G**) Representative confocal images of WT and C9orf72 KO microglia that have engulfed fluorescent beads (red), stained by IB4 (green), and DAPI (grey). Scale bar 20 µm. Right, quantification. Each data point represents mean percentage of phagocytic microglia per experiment: n=30 fields/genotype from N=3. Unpaired t test, p=0.7285. **H**) Representative confocal images of WT and C9orf72 KO microglia stained by IB4 (green) and DAPI (grey) at 4 hours after incubation with pHrodo-synaptosomes (red) (T0) and at 4 hours after washout (T4). Scale bar 20 µm. Right, quantification of the IB4+ area covered by synaptosomes. Data points represent mean microglial area covered by synaptosomes in C9orf72 KO microglia normalized on WT per experiment: n≥40 analyzed fields per genotype from N=3. Mann Whitney test: WT T0 vs. KO T0 p=0.6857, Unpaired t test: WT T4 vs. KO T4 p=0.0441.

### C9orf72 KO microglia engulf more synaptic terminals *in vitro*

Next, we co-cultured neurons with WT or C9orf72 KO microglia and quantified synaptic density and microglial synaptic engulfment. The density of Bassoon+ puncta in neurons co-cultured with C9orf72 KO microglia, releasing more EVs, was lower compared to neurons co-cultured with WT microglia or cultured alone (Fig. 4A-B), strengthening the link between EVs production and synaptic loss. Accordingly, analysis of Bassoon+ and Shank2+ material inside microglia, revealed higher uptake of synaptic structures in C9orf72 KO microglia compared to WT cells (Fig. 4C-E). Because the phagocytic capacity of C9orf72 KO microglia is equal to that of WT cells (Fig. 3G-H), we reasoned that lower pre-synaptic density and higher synaptic engulfment by C9orf72 KO microglia may be the consequence of higher EVs-mediated C1q secretion and deposition to synapses. Thus, we asked whether partial inhibition of EVs release from C9orf72 KO microglia could rescue normal pre-synaptic density.

**Figure 4.**
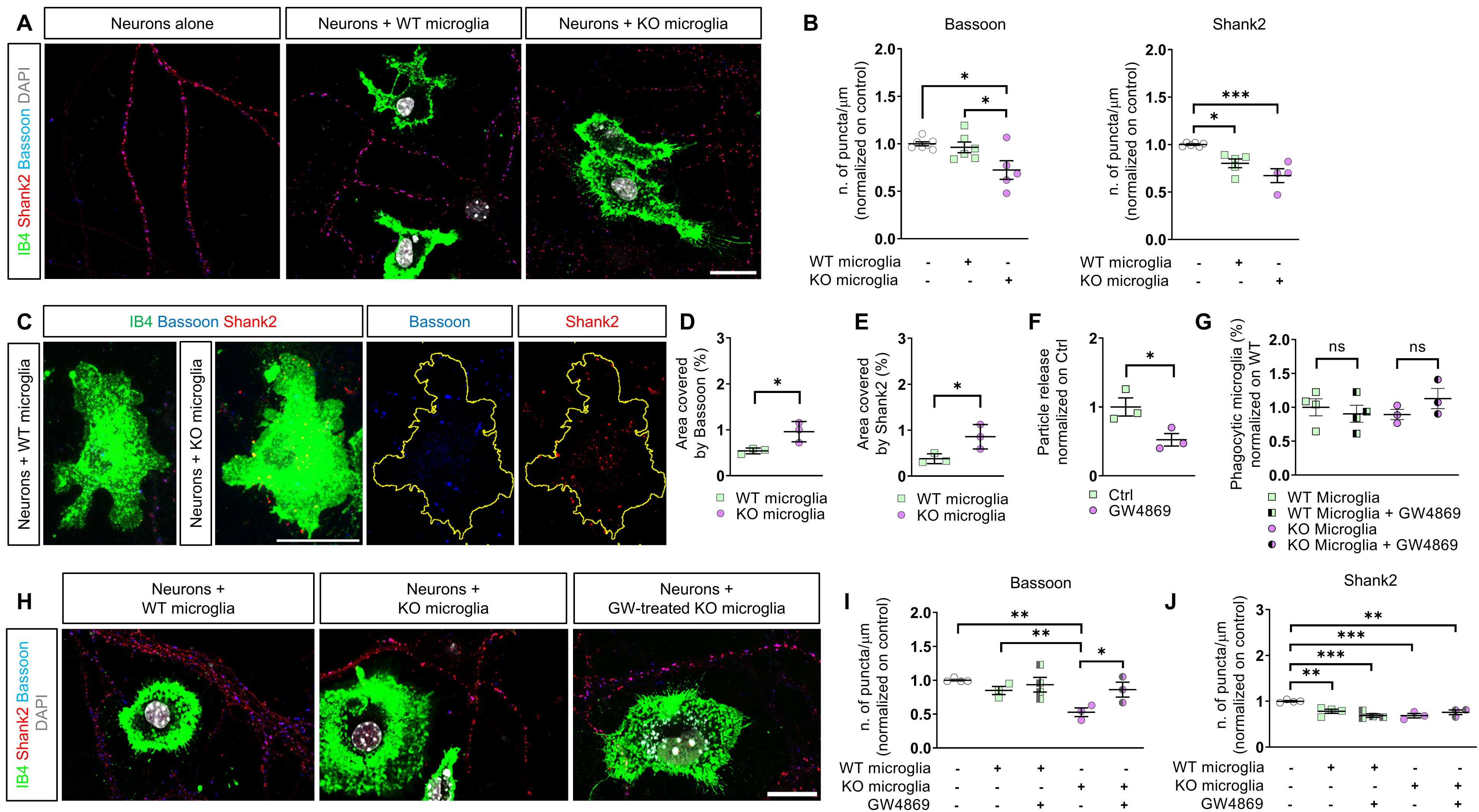
C9orf72 KO microglia engulf more synaptic material compared to WT cells. **A**) Representative confocal acquisition of neurons cultured alone, with WT or C9orf72 KO microglia, stained by IB4 (green) and for Shank-2 (red), Bassoon (blue) and nuclei (grey). Scale 20 µm. **B**) Relative quantification of pre- and post-synaptic density. Data points represent mean synaptic density along dendrites normalized on neurons alone per experiment. Pre-synaptic density: n≥150 dendrites per condition from N≥5. One-way ANOVA with Holm-Sidak’s multiple comparisons test: neurons alone vs. KO microglia p=0.0146, WT microglia vs. KO microglia p=0.0290. Postsynaptic density: n≥120 dendrites per condition from N≥4. Neurons alone vs. Neurons with WT microglia p=0.0102, Neurons alone vs. Neurons with KO microglia p=0.0006, Neurons with WT microglia vs. with KO microglia p=0.0653. **C**) Representative z-stack projections of IB4+ WT and C9orf72 KO microglia (green) cocultured with neurons, stained for Bassoon (blue) and Shank2 (red). Right images show Bassoon and Shank2 puncta inside the perimeter (yellow line) of KO microglia. Scale bar 20 µm. **D-E**) Quantification of Bassoon+ (**D**) and Shank2+ (**E**) material engulfed by WT and KO microglia. Data points represent the percentage of IB4+ area stained by Bassoon/Shank2 per experiment: n=45 analyzed fields per genotype from N=3. Unpaired t test: Bassoon p=0.0345, Shank2 p=0.0447. **F**) Constitutive EVs production from microglia at 24 hours following GW4869 treatment, analysed by TRPS. Data points represent single measurements normalized to control from N=3. Unpaired t test p=0.0409. **G**) Percentage of WT and C9orf72 KO microglia treated or not with GW4869, which have engulfed fluorescent beads. Data points represent the percentage of phagocytic microglia normalized on untreated WT microglia per experiment: n≥30 analyzed fields per condition from N=3. One-way ANOVA with Holm-Sidak’s multiple comparisons test: p>0.8027. **H**) Representative confocal images of neurons cultured with IB4+ WT or C9orf72 KO microglia pre-treated or not with GW4869, stained for Shank2 (red), Bassoon (blue) and nuclei (grey). Scale bar 20 µm. **I-J**) Quantification of pre- and post-synaptic density. Data points represent synaptic density normalized to neurons alone per experiment: n≥30 analysed dendrites/condition from N≥3. Bassoon: One-way ANOVA with Fisher’s LSD test: neurons alone vs. with KO microglia p=0.0013, neurons with WT vs. with KO microglia p=0.0213, neurons with KO vs. with GW4869-treated KO microglia p=0.0177. Shank2: One-way ANOVA with Holm-Sidak’s multiple comparisons test: neurons alone vs. with WT microglia p=0.0091, vs. with GW4869-treated WT microglia p=0.0005, vs. with KO microglia p=0.0009, vs. with GW4869-treated KO cells p=0.0080.

We incubated overnight C9orf72 KO microglia (and WT cells) with GW4869 (5 μM), a known inhibitor of constitutive EVs release (Trajkovic et al., 2008), and found that EVs production was reduced by ∼50% at 24 hours after treatment, as evidenced by TRPS measurements (Fig. 4F). Thus, we co-cultured GW4869-treated microglia with neurons for 24 hours and analysed synaptic density. Quantification of Bassoon+ and Shank2+ puncta in dendrites closed to microglia revealed that GW4869 treatment restored normal pre-synaptic density in neurons cocultured with C9orf72 KO microglia while left the density of Shank2 post-synaptic puncta unaltered (Fig. 4H-J). Importantly, treatment with GW4869 did not influence the phagocytic capacity of microglia, as measured by fluorescent beads internalization (Fig. 4G). No significant changes in the density of pre- and post-synaptic puncta were observed in neurons cocultured with GW4869-treated WT microglia (Fig. 4I-J).

Taken together these data indicated that EVs production controls synaptic pruning, reflecting the ability of EVs to tag pre-synaptic terminals with C1q.

### Larger production of C1q positive EVs from C9orf72 KO microglia in the adult C9orf72 KO brain

C9orf72 deficiency in microglia was previously associated with enhanced complement-mediated microglial synaptic removal in adult mice (Lall et al., 2021). Thus, we examined microglial EVs production and EVs-mediated C1q secretion from C9orf72 KO and WT adult brain.

We adapted previous protocols (Crescitelli et al., 2021; D’Acunzo et al., 2022; Vella et al., 2017) to optimize extraction of EVs from the interstitial fluid of brain tissue (cortex and hippocampus) through a mild extracellular matrix digestion followed by a series of filtrations and centrifugations steps. Two crude EVs fractions were collected by centrifugation at *10000x g* (low speed EVs, l-EVs) and *120000x g* (high speed EVs, h-EVs), which were further purified on iodixanol density step gradient (Fig. 5A).

**Figure 5.**
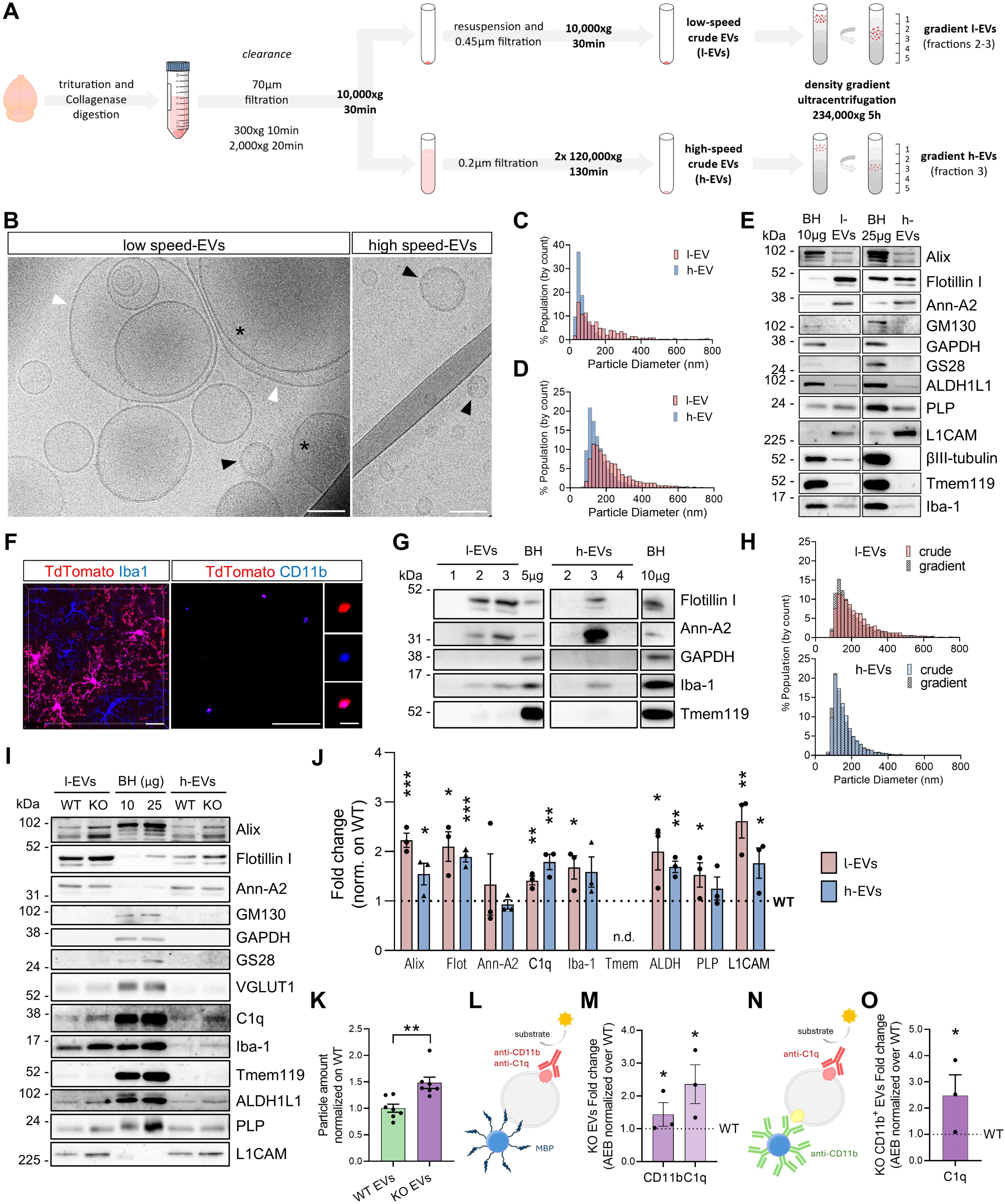
Higher EVs production from C9orf72 KO microglia in the mouse brain. **A**) Schematic of EVs extraction protocol from mouse brain. **B**) Representative cryo-EM micrographs of crude low speed EVs (l-EVs) and high speed EVs (h-EVs). White arrowheads point to multiple-membrane bound EVs or EVs with intraluminal vesicles; black arrowheads to EVs with rough surface; asterisks to electron dense EVs. Scale bars 100nm. **C-D**) Size distribution of l-EVs and h-EVs, detected by cryo-EM (C) and TRPS (D). **E**) Representative WB of l-EVs and h-EVs from adult mouse brain (50 mg, 4.5 months old) for EVs markers, intracellular and brain cell lineage markers. Brain homogenate (BH) is loaded as a positive control for contaminants. **F**) Imaris 3D projection of the hippocampal region of a Cdh5-CreER^T2^::R26^tdTomato^ transgenic mouse brain in which Cre recombination was induced with one single pulse of 4-hydroxytamoxifen at E7.5 expressing tdTomato (red) in microglia, stained for Iba-1 (blue) (left). Scale bar 20 µm. Representative confocal image of EVs from the transgenic brain stained for CD11b (blue) (right). Scale bar 10 µm, Zoom-in scale bar 2 µm. **G**) Representative WB of purified l-EVs (fractions 1,2,3) and h-EVs (fractions 2,3,4) for EVs positive and negative markers, Iba-1 and Tmem119. Fractions densities (g/ml): 1= 0.06, 2= 0.07-0.09, 3= 0.12-0.13, 4= 1.21-1.22. **H**) Size distribution of crude vs purified brain EVs fractions (l-EVs and h-EVs). **I**) WB of crude l-EVs and h-EVs from an equal amount of WT and C9orf72 KO brain (50 mg) stained for EVs markers, the synaptic vesicles marker VGLUT1, cell markers and C1q (**J**) Relative densitometric quantification. Data points represent KO measurements normalized on WT from n=3 mice/genotype. T-test): Alix l-EVs p=0.0005, h-EVs p=0.0334; Flotillin I l-EVs p=0.0106, h-EVs p=0.0005; Annexin A2 ns; C1q l-EVs p=0.0046, h-EVs p=0.0035; Iba-1 l-EVs p=0.0223, h-EVs ns; ALDHL1 l-EVs p=0.0272, h-EVs p=0.0018; PLP l-EVs p<0.050, h-EVs ns; L1CAM l-EVs p=0.0046, h-EVs p=0.0332. **K**) Quantification of crude l-EVs and h-EVs pooled together from WT and C9orf72 KO brains analysed by TRPS. Data points represent single measurements from n=3 mice/genotype, normalized on WT cells. Mann-Whitney test p=0.002. **L-M**) Schematic of the assay (**L**) and SiMoA quantification of CD11b+ and C1q+ l-EVs from WT and C9orf72 KO brains (n=3 mice/genotype) (**M**). AEB (Average Enzyme per Bead) signal was obtained using MSP modified beads and CD11b or C1q antibody as detector. Marker levels of C9orf72 KO EVs were normalized on WT. Mann-Whitney test: CD11b and C1q p= 0.0381. **N-O**) Schematic of the assay using CD11b conjugated beads and C1q antibody as detector (**N**) and SiMoA quantification of C1q surface cargo in CD11b+ EVs from WT and C9orf72 KO brains (n=3 mice/genotype) (**O**).

The purity, quantity and cell source of crude EVs fractions were characterized by cryo-EM, TRPS, WB and confocal microscopy. According to Cryo-EM, both l-EVs and h-EVs consisted of membrane EVs round in shape, heterogeneous in size (from 11 to 890 nm) and content of electron-dense material, some of which were multiple-membrane bound or contained intraluminal vesicles (Fig. 5B, white arrowheads) or had rough surface (Fig. 5B, black arrowheads). l-EVs were larger compared to h-EVs, as evidenced by both cryo-EM (n=341 l-EVs, n=590 h-EVs; Mann-Whitney test p<0.0001; Fig. 5C) and TRPS analysis (Mann-Whitney test p=0.0159; Fig. 5D). WB analysis showed that both l-EVs and h-EVs were enriched in the EVs markers Flotillin I and Annexin-A2 compared to brain homogenates and depleted in intracellular markers (GM130 and GS28 for Golgi, GAPDH for cytosol, VGLUT1 for synaptic vesicles), excluding gross contamination with intracellular organelles (Fig. 5E, 5I and S3, S6 for protein loading). As expected, brain EVs stained positive for oligodendroglia (PLP), astrocyte (ALDH1L1), and neuron (L1CAM, βIII-tubulin) lineage markers, and for the microglia and brain macrophage marker Iba-1. The latter was more abundant in l-EVs compared to h-EVs, despite the higher amount of loaded h-EVs (Fig. 5E, S3 for protein loading). This is in line with accumulation of Iba-1 in large EVs budding *in vitro* from the microglial surface, which are mostly recovered at low centrifugation speed (Verderio et al., 2012). Conversely, the microglial lineage marker Tmem119 was largely depleted in both brain EVs fractions compared to brain homogenate and barely detectable by WB analysis (Fig. 5E, I), hindering the detection of EVs uniquely derived from microglia.

Microglia originate during development from yolk sac (YS)-derived erythromyeloid progenitors (Kierdorf et al., 2013). Previous fate mapping experiments showed that Cdh5-CreER^T2^::R26^tdTomato^ transgenic mice (Azzoni et al., 2014), which inducibly express tdTomato fluorescent protein under the control of VE-Cadherin (Cdh5), can reliably be used to trace the progeny of YS hemogenic endothelium, including microglia (Gentek et al., 2018). Thus, to show the presence of microglia-derived EVs *in vivo*, we next extracted EVs from the brain of Cdh5-CreER^T2^::R26^tdTomato^ mice. Flow cytometric analysis of the transgenic brains showed that a single dose of 4-hydroxytamoxifen at embryonic day (E)7.5 resulted in tdTomato labelling of more than 90% of TMEM119+ microglia and only a few CD31+ endothelial cells, (Fig. S4) consistent with previous data (Gentek et al., 2018). EVs isolated from Cdh5-CreER^T2^::R26^tdTomato^ brains were stained for the microglial/macrophage marker CD11b and imaged by confocal microscopy. EVs co-expressing td-Tomato+ and CD11b+ EVs were detected, unequivocally demonstrating the microglial origin of a subpopulation of brain EVs (Fig. 5F). Further purification of crude EVs on iodixanol density step gradient revealed a floating density of 1.07-1.13 g/ml for l-EVs and of 1.12-1.13 g/ml for h-EVs, both within the known range of EVs (D’Acunzo et al., 2022) (Fig. 5G and S5 for protein loading) and confirmed larger Iba-1 expression relative to Tmem119 in purified EVs (Fig. 5G). According to TRPS measurements, purified and crude EVs had the same size distribution (Fig. 5H), supporting the absence of contaminants in crude preparations.

Given that crude EVs fractions were devoid of gross contamination while assuring rapidity of the procedure and higher yield, crude EVs fractions were employed to measure EVs production from C9orf72 KO and WT microglia/macrophages in the adult brain, using the myeloid marker Iba-1. Iba-1 was overrepresented in l-EVs and h-EVs (by 1.68 and 1.58-fold, respectively) from C9orf72 KO brain compared to WT, along the EVs marker Flotillin I and Alix (Fig. 5I-J and S6 for protein loading, n=3 mice/group), suggesting augmented production of total and microglial EVs in the mutant brain. ALDH1L1, PLP and L1CAM were also more abundant in EVs from C9orf72 KO brain compared to WT, suggesting a general activation of C9orf72 KO brain cells. Staining of brain EVs for C1q, to examine EVs-mediated C1q secretion, showed low levels of the complement factor in EVs compared to brain homogenate (Fig. 5I), suggesting low production of C1q carrying EVs in the adult mouse brain. However, C1q was upregulated in EVs samples from C9orf72 KO brains compared to WT (Fig. 5I-J), revealing augmented release of C1q+ EVs in the mutant brain. Crude EVs (l-EVs and h-EVs together) from C9orf72 KO and WT brain were further quantified using TRPS analysis which confirmed higher EVs production in the mutant brain (Fig. 5K, n=3 mice/genotype). In addition, we further estimated the relative proportion of C1q+ EVs and EVs derived from C9orf72 KO and WT microglia/macrophages by Single Molecule Array (SiMoA) technology. EVs were captured by using paramagnetic beads functionalized with a membrane sensing peptide (MSP) to act as a pan-vesicular binding probe, as previously reported (Gori et al., 2024). EVs were subsequently probed with antibodies against C1q and CD11b, confirming higher abundance of both EV populations in the mutant brain (Fig. 5 L-M). Finally, by SiMoA, we further measured C1q content in EVs derived from WT and mutant microglia/macrophages. In this case, EVs were captured by beads functionalized with anti-CD11b and probed with anti-C1q antibodies (Fig. 5N). The assay showed higher signal detection for C1q in CD11b+ EVs from C9orf72 KO brain (Fig. 5O). Collectively, these data confirmed in vivo increased production of C1q carrying EVs from C9orf72 KO microglia, and revealed a general increase in EVs production from C9orf72 KO brain cells.

### C9orf72 KO microglia engulf more synaptic terminals in the early postnatal hippocampus

Next, we asked whether larger EVs production from C9orf72 KO microglia might influence microglia-mediated synaptic pruning during brain development. To address this question, we analysed the CA1 hippocampal region from WT and C9orf72 KO mice at postnatal day (P)17, a time of high synapse remodelling. Quantification of Iba-1+ cells revealed no changes in microglial density in C9orf72 KO mice compared to WT (Fig. 6A) whereas three-dimensional (3D) microglial reconstruction confirmed accumulation of lysosomal proteins in C9orf72 KO microglia, as evidenced by larger volume of CD68+ structures per individual cell (Fig. 6B).

**Figure 6.**
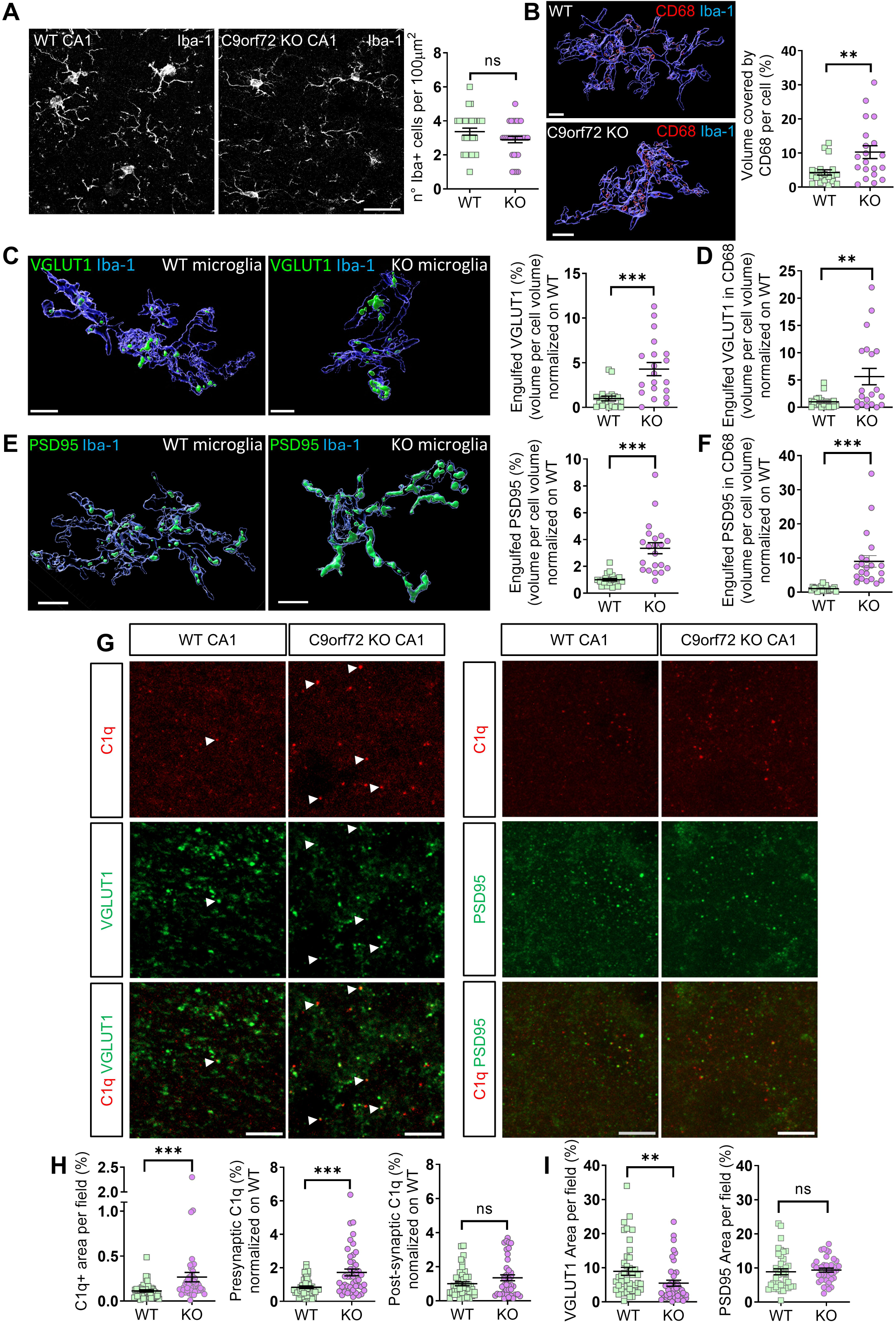
C9orf72 KO microglia remove more synaptic terminals in the hippocampus at P17. **A**) Representative z-projections of CA1 from WT and C9orf72 KO mice stained for Iba-1 (grey) at P17. Scale bar 20 µm. Right: relative quantification of microglial density. Each data point represents microglia density per field, n=30 fields from n=3 mice/genotype. Unpaired t test, p=0.0997. **B**) Representative 3D reconstruction of microglia stained for Iba-1+ (blue) and CD68 (red) from WT and KO CA1 at P17, scale bar 5 µm. Right, corresponding volume of CD68+ structures. Data points represent single cells, n=20 cells from n=3 mice/genotype Mann-Whitney test, p=0.0056. **C**) Representative 3D reconstruction of Iba-1+ microglia (blue) engulfing VGLUT1+ puncta (green) from WT and KO CA1 regions. Scale bar 5 µm. Right: quantification of engulfed VGLUT1 per microglia, n=20 cells from n=3 mice/genotype. Each data point represents the percentage of VGLUT1+ over Iba-1+ volume (Mann-Whitney test p<0.0001). **D**) Quantification of VGLUT1 in CD68+ endolysosomes. Data points represent the percentage of VGLUT1 and CD68 double+ volume over Iba-1+ volume (Mann-Whitney test p=0.0095). **E**) 3D reconstruction of Iba-1+ microglia (blue) engulfing PSD95+ puncta (green) from WT and KO CA1 regions. Scale bar 5 µm. Right: quantification of engulfed PSD95 per microglial cell, n=21 cells from n=3 mice/genotype. Each data point represents the percentage of PSD95+ over Iba-1+ volume (Mann-Whitney test p<0.0001). **F**) Quantification of engulfed PSD95 in the CD68+ compartment. Each data point represents the percentage of PSD95 and CD68 double+ volume over Iba-1+ volume (Mann-Whitney test p<0.0001). **G**) Representative z projections of CA1 from WT and KO mice double stained for C1q and VGLUT1/PSD95 at P17. White arrowheads indicate some double positive puncta. Scale bar 5 µm. **H**) Quantifications of the percentage of C1q+ area per field (left), n=47 fields from n=3 mice/genotype (Mann-Whitney test p<0.0001), of the percentage of C1q+ VGLUT1+ terminals per field normalized on WT (middle), n=45 fields from n=3 mice/genotype (Mann-Whitney test p=0.0002) or C1q+ PSD95+ terminals per field normalized on WT (right), n≥41 fields, from n=3 mice/genotype (Mann-Whitney test p=0.2383). **I**) Quantification of pre- and post-synaptic density. Data points represent the percentage of VGLUT1+ area per field, n=42 fields (Mann-Whitney test p=0.0028) or PSD95+ area per field, n≥38 fields (Mann-Whitney test p=0.2501) from n=3 mice/genotype.

Further analysis revealed that microglia from C9orf72 KO mice contained significantly more synaptic material, VGLUT1+ and PSD95+, in the cytoplasm and within CD68+ phagolysosomes, when compared to WT (Fig. 6C-F), indicating enhanced engulfment. However, analysis of C1q immunoreactivity in the CA1 region showed higher C1q deposition only on pre-synaptic VGLUT1+ but not PSD95+ puncta in C9orf72 KO mice (Fig. 6G-H), suggesting that EVs-mediated C1q tagging occurred on the pre-synaptic compartment. Moreover, analysis of synaptic density revealed lower density only for pre- but not post-synaptic terminals in C9orf72 KO mice (Fig. 6G, I), despite increased pre- and post-synaptic engulfment by C9orf72 KO microglia. This suggested that presynaptic removal may trigger postsynaptic pruning, leading to postsynaptic loss at later postnatal stages. Collectively these data linked augmented microglial EVs production from C9orf72 KO microglia to enhanced pre-synaptic C1q deposition and microglia-mediated synaptic pruning in the hippocampal region during the critical period of circuit refinements.

### Microglia release more EVs during the pruning period in the postnatal mouse brain

The link between microglial EVs production and synaptic pruning established in C9orf72 KO brain prompted us to ask whether microglia may release more EVs during physiological pruning across development.

We first interrogated the Microglia Development RNASeq searchable database (https://microglia-seq.vm.duke.edu/microglia-seq/shiny-app/) (Hanamsagar et al., 2017) to investigate whether known EVs markers and factors mediating EVs release may be upregulated in mouse microglia across brain development. The analysis revealed upregulation of a few genes enriched in EVs (Alix, AnxA1, AnxA5) or involved in EVs biogenesis/release (Arf6, Arrdc1, Vps37b, Sepp1, rab27b) at time points critical for synaptic pruning during postnatal development, i.e at the onset of synaptic pruning in the dorsolateral geniculate nucleus (dLGN) of the thalamus (P4-P5), and/or during the pruning period in the hippocampus (P14), compared to periods preceding (E14 and E18) or following (P60) the pruning window (Fig. S7). These results supported a possible increase in microglial EVs biogenesis and release during developmental pruning.

Then we measured by WB the yield of microglial EVs extracted from the mouse brain (deprived of olfactory bulbs, cerebellum, pons and medulla but including thalamus with dLGN, Fig. S8A) at P2, before the onset of synaptic pruning, at P7 and P17, during the peak pruning period in the dLGN and the hippocampus respectively and in the adult brain, after developmental synaptic pruning (Soteros and Sia, 2022). The analysis revealed higher Iba-1 expression in l-EVs at P17 compared to P2, P7 and the adult brain, suggesting higher production of microglia/macrophage-derived EVs during the pruning activity in the hippocampus (Fig. 7A-C and S8B-C for protein loading, n=3-7 mice/time-point).

**Figure 7.**
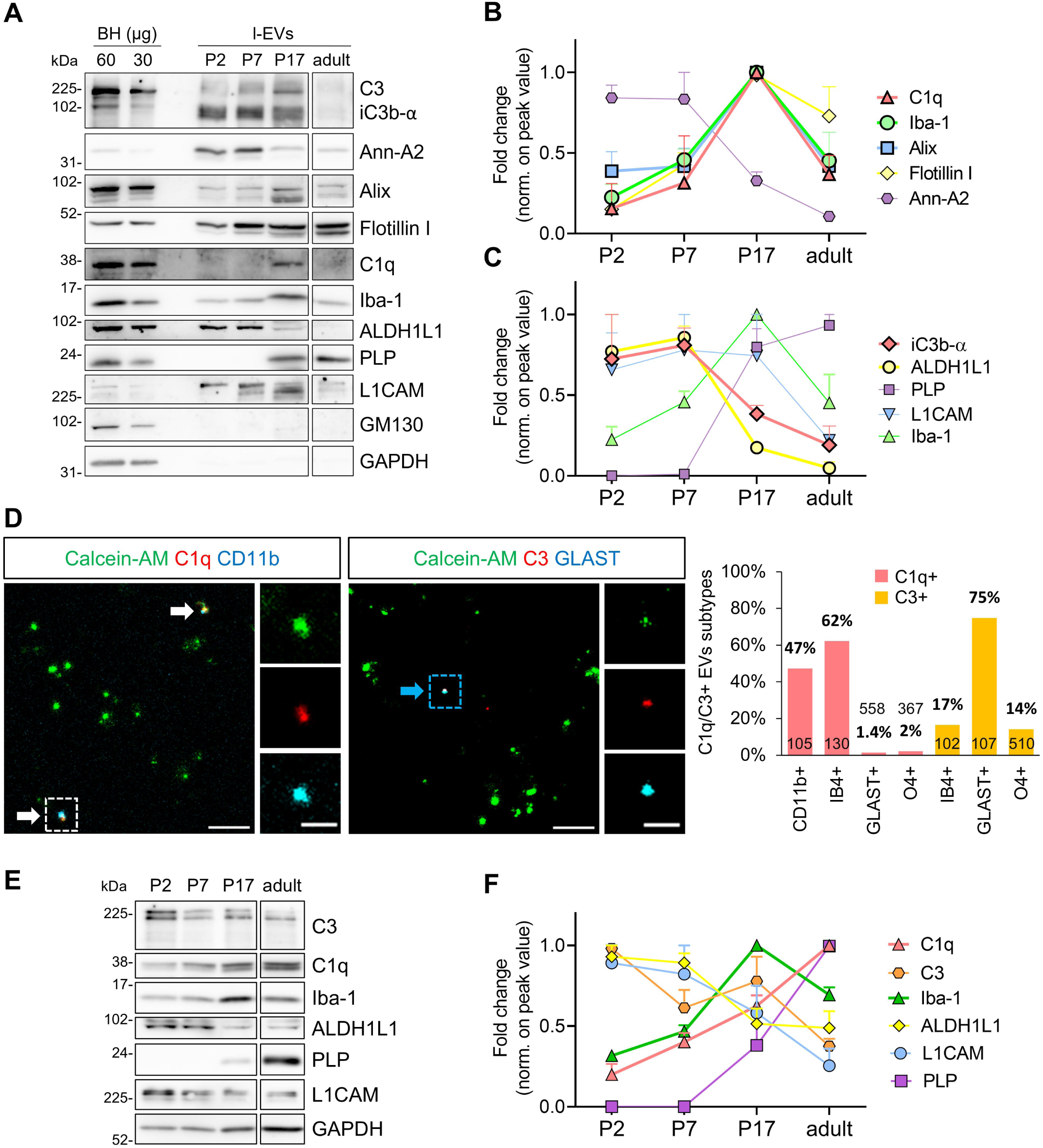
Microglial EVs secretion across postnatal brain development. **A**) Representative WB of l-EVs extracted from an equal amount of WT brain (75 mg) at P2, P7, P17 and adult age stained for EVs makers, C1q, C3, Iba-1 and other cell lineage markers. **B-C**) Corresponding densitometric quantification of C1q, Iba-1 and EVs markers (**B**), or iC3b fragment and cellular markers (**C**). For each marker, data are normalized on peak value. Data are from a total of 7 P2, 4 P7, 3 P17, 3 adult brains (N=3 independent extractions). **D**) Representative fluorescence confocal images of brain l-EVs extracted at P17 and stained for C1q (red), CD11b (light blue) and calcein-AM (green) (left) or for C3 (red), GLAST (light-blue) and calcein-AM (green) (middle). CD11b+ EVs are indicated by white arrows, GLAST+ EVs by light blue arrow. Scale bar 5 µm. Zoom-in of dotted squares are on the right. Scale bar 2 µm. Quantification of C1q+/C3+ EVs subtypes, immunoreative for the cell lineage markers CD11b, IB4, GLAST or O4. Number of analysed EVs are indicated for each EV population. **E-F**) Representative WB of brain homogenates (40 μg) from P2, P7, P17 and adult mice, stained for cellular markers, complement factors, and GAPDH as loading control (**E**), and relative densitometric quantification (**F**). For each marker, data are normalized on peak value (n=3 mice/time-point).

Of note, l-EVs immunoreactivity for C1q and Alix overlapped Iba-1 staining at all time points. Conversely, Annexin-A2, iC3b α-chain fragment and the astrocytic marker ALDH1L1 were largely expressed at P2-P7 (Fig. 7A-C). WB analysis of h-EVs confirmed the peak of Iba-1 expression at P17, and higher iC3b α-chain fragment, ALDH1L1 and Annexin A2 immunoreactivity at P2 and P7, while C1q staining was barely detectable (Fig. S11). Taken together these findings suggested that Alix positive microglia-derived EVs, enriched in the l-EVs fraction, may act as main carriers for released C1q during high synapse remodeling in the hippocampus (P17), whereas astrocytes-derived EVs might represent the primary delivery vehicles for secreted iC3b at earlier postnatal developmental stages.

To definitely demonstrate the microglial origin of C1q-carrying EVs produced during the pruning period in the hippocampus, we next performed surface immunofluorescence staining of l-EVs for calcein-AM, an index of EVs integrity, the microglia/macrophage markers CD11b-Cy5 or IB4-568 and C1q at P17, the time point of largest C1q secretion across development. About 47% of CD11b+ l-EVs and 62% of IB4+ l-EVs were immunoreactive for C1q (Fig. 7D) whereas a very low percentage of EVs from astrocytes (GLAST+) or oligodendrocytes (O4+) stained positive for C1q (1.4% and 2% respectively). Conversely about 75% of GLAST+ EVs, 17% of IB4+ EVs and 14% of O4+EVs were immunoreactive for C3 (Fig. 7D), indicating astrocytes-derived EVs as main carriers for secreted C3.

Finally, we asked whether increased yield of Iba-1 EVs at P17 reflected developmental microglia expansion or larger microglial EVs production. WB analysis of brain homogenates across development revealed highest Iba-1 expression at P17, indicative of peak microglial expansion (Fig. 7E-F and S10 for Ponceau staining; n=3 mice/time-point). This suggested that higher yield of Iba-1+ EVs at the time point of high synaptic remodelling in the hippocampus likely reflects microglial expansion. Of note, WB analysis revealed that C1q protein levels progressively increased across postnatal development, peaking in the adult brain (Fig. 7E-F), when the amount of C1q+ EVs was very low (Fig. 5I and 7A). This finding is in line with the well-known increase in C1q protein content during brain aging (Stephan et al., 2013) and recent evidence that microglial secreted C1q is localized inside neurons in the adult and aging brain, where it alters neuronal protein translation (Scott-Hewitt et al., 2024).

## DISCUSSION

Several studies have indicated that the complement factor C1q is released from microglia during brain development (Fonseca et al., 2017), localizes to inappropriate synapses and acts as a tag for synaptic pruning (Schafer et al., 2012; Stephan et al., 2012; Stevens et al., 2007). More recently, it has been clarified that binding of C1q to weak synapses is guided by PS, a neuronal eat-me signal (Li et al., 2020; Scott-Hewitt et al., 2020), which is focally externalized on synapses following downregulation of the phospholipid-flippase chaperone CDC50A upon neuronal inactivation (Li et al., 2021). However, how C1q reaches PS on synapses in need to be removed remains obscure.

Here we show that C1q is secreted from microglia and delivered to pre-synapses by means of EVs, and provide a mechanistic link between EVs production by microglia and microglial synaptic removal. *In vitro,* we show that microglial EVs i) are highly enriched in C1q compared to donor microglia, as indicated by WB, and carry 60% of secreted C1q, according to ELISA quantification, ii) make contacts with synapses, as evidenced by both live imaging of individual EVs on mature dendrites and confocal analysis of neurons exposed in bulk to fluorescent EVs, iii) selectively mark PS positive pre-synapses with C1q, as indicated by increased pre-synaptic C1q deposition in neurons supplemented with EVs but not in annexin V-treated neurons, in which PS residues are masked, iv) promote synaptic loss upon supplementation to neurons or when released in higher amount by mutant (C9orf72 KO) microglia, whereas normalization of EVs production rescues normal pre-synaptic density in co-cultured neurons. *In vivo,* we show i) higher production of both total EVs and microglial EVs carrying C1q in C9orf72 KO brain compared to WT, as indicated by TRPS, SiMoA quantification and WB analysis and using EVs markers (Flotillin, Alix) microglia cell lineage markers (Iba-1, CD11b) and anti-C1q antibody, and ii) augmented C1q deposition at pre-synaptic sites in the hippocampus of KO mice during the pruning period, resulting in lower pre-synaptic density and larger microglial synaptic engulfment. Finally, we show that in postnatal mouse brains production of microglial EVs carrying C1q correlates with peak pruning activity in the hippocampus.

Overall, our data support a model in which microglia package C1q into EVs for precise deposition of the immune factor to inappropriate synapses and release more EVs to promote removal of targeted synaptic terminals during active pruning period.

### C1q and C3 are released by microglia as components of EVs

Complement factors are traditionally known as soluble molecules produced locally by immune-cells and activated extracellularly to engage their receptors on adjacent cells and induce specific responses (West and Kemper, 2023). However, consolidated evidence indicates that complements are released by immune cells as surface cargo of EVs, generated either inside multivesicular bodies (exosomes) or at the cell surface (ectosomes) (Karasu et al., 2018).

Already twenty years ago macrophages and immature dendritic cells (Castellano et al., 2004), important cell sources of C1q in the periphery, were shown to express a membrane-bound form of C1q, thus being able to shed C1q-bearing EVs directly from the plasma membrane. Furthermore, macrophages and B cells have long been known to release exosomes carrying C3 as a mechanism of elimination of C3 fragments deposited on the cell membrane (Papp et al., 2008). The underlying mechanism involves C3 fragments endocytosis, followed by their targeting to multivesicular bodies (MVBs), binding to the surface of intraluminal vesicles (pre-secreted exosomes) and secretion as exosome surface components upon MVBs fusion with the plasma membrane (Buzas et al., 2018). Recently, it has been reported that complement factors can also associate with the outside of EVs once the EVs are released by the producing cell in extracellular fluids, forming a protein corona around EVs with other surface molecules, including extracellular matrix components, cytokines and enzyme*s* (Buzas, 2022).

In line with these findings, we hereby show that cultured microglia release C1q in association with EVs and that C1q at EVs surface accounts for most of the complement content in microglial supernatant. Moreover, by SIMOA or immunofluorescence staining, we validate the presence of C1q on the surface of CD11b+ EVs produced by microglia/macrophages in the adult mouse brain and at peak vesicular C1q secretion across postnatal development (P17). C1q association with the EVs membrane in the endocytic compartment or outside the cells can be mediated by PS, which binds the globular head of C1q avidly (Paidassi et al., 2008) and is externalized on microglia-derived EVs, especially on large ectosomes budding from the cell surface (Bianco et al., 2009; Buzas, 2022). Finally, by confocal analysis we identify GLAST+ astrocytes-derived EVs as main carrier for C3/iC3b, and show that the production of C3-storing EVs is particularly elevated at very early postnatal stages, when the release of C1q-bearing EVs is still low, as indicated by the temporal plot of brain EVs marker expression across development. In line with these findings, abundant localization of small astrocytes-derived EVs has been previously reported outside of astrocytes in rodent brain during the first postnatal week (Jin et al., 2023). Further work is needed to explore whether at very early postnatal stages C3-bearing EVs from astrocytes might contribute to microglial synaptic engulfment in specific brain regions through an alternative complement cascade, which is independent of C1q localization to synapses and works through direct localization of C3 to dysfunctional synapses (Werneburg et al., 2020).

### What are the advantages of EVs-mediated complement secretion and deposition to the synapse?

First, being released as surface cargo, C1q and other complement factors are protected from dilution and can efficiently engage their ligands, such as externalized PS, on synapses.

Second, by moving at the neuron surface and scanning the plasma membrane (D’Arrigo et al., 2021; Gabrielli et al., 2022), EVs can precisely bring surface C1q in contact with synapses that externalized PS, favouring its deposition on weak synapses. In line, pre-synaptic terminals, the main synaptic targets of microglial EVs according to this study, are the sites of preferential PS externalization during the pruning period in the developing brain (Scott-Hewitt et al., 2020). In this scenario, PS residues externalized on weak synapses are expected to compete with those present at the EVs surface for binding to C1q, while it is tempting to hypothesize that other C1q ligands, among multiple receptors and multimolecular complexes co-expressed with PS on synapses (Thielens et al., 2017), and /or larger amount of externalized PS may facilitate the transfer of the immune factor from microglial EVs to pre-synaptic sites (Fig. 1K)

Third, EVs-associated complement factors can signal to distant brain regions and reach the blood. Brain-to-blood transit of EVs containing complements has been demonstrated for astrocytes-derived EVs and analysis of their cargoes proven to capture complements alterations not detectable in total plasma (Burgelman et al., 2022), helping the diagnosis of Alzheimer’s disease and the prediction of mild cognitive impairment conversion to disease (Goetzl et al., 2018; Winston et al., 2019).

Interestingly, we recently identified the “spare me” signal CD47 (Lehrman et al., 2018) in the proteome of EVs released by primary microglia, in addition to complement factors (Gabrielli M., Verderio C., unpublished proteomic data). This suggests that microglial EVs may act as carriers of both “eat me” and “don’t eat me” signals, being able to tag not only weak synapses for removal but also active synapse for protection from excessive pruning. Further investigation is required to test this intriguing hypothesis and clarify whether C1q and CD47 are carried on the same or distinct EVs produced by microglia.

### EVs-mediated synaptic pruning occurs via selective C1q deposition at pre-synaptic terminals

A key finding of this study is the observation that microglial EVs preferentially drive pre-synaptic loss *in vitro* and *in vivo* and that pre-synaptic loss requires a threshold amount of microglial EVs to occur. This amount is provided by C9orf72 KO microglia or can be reached by EVs supplementation to neurons cultured with WT cells. Specifically, we show that larger EVs production from C9orf72 KO microglia results in higher loss of pre-synaptic compared to post-synaptic sites in microglia-neurons cultures and decreased pre-but not post-synaptic density in the hippocampus of C9orf72 KO mice at P17, during pruning period. Conversely, pharmacological inhibition of EVs release from C9orf72 KO (and WT) microglia does not influence the density of post-synaptic terminals while prevents pre-synaptic loss in neurons co-cultured with mutant microglia. In addition, EVs supplementation to neurons specifically increases C1q deposition to pre-synaptic sites and reduces the density of pre-synaptic but not post-synaptic sites in neurons cultured with WT microglia. Taken together, these data are consistent with previous studies showing that pre-synaptic microglial engulfment is a complement (and CR3 dependent) process (Schafer et al., 2012; Scott-Hewitt et al., 2020) while suggest that postsynaptic removal may occur in a complement independent manner, e.g. via phagocytic receptors like TREM-2, BAI1 or GPR56 that bind externalized PS directly, as previously reported (Scott-Hewitt et al., 2020). However, it should be highlighted that the loss of pre-synaptic terminals and pre-synaptic microglial engulfment detected under elevated microglial EVs concentration were accompanied by increased uptake of post-synaptic material by microglia *in vitro* and in the C9orf72 KO hippocampus, suggesting that EVs-mediated C1q pre-synaptic deposition and pre-synaptic removal may indirectly engage post-synaptic engulfment. This may lead to postsynaptic loss at later postnatal stages, contributing to the synaptic dysfunction due to C9orf72 haploinsufficiency (Bauer et al., 2022; Lall et al., 2021). Further studies are warranted to elucidate these aspects.

### Production of microglial EVs increases during the pruning period in the postnatal brain

A key unanswered question related to complement-mediated pre-synaptic removal is about the mechanism(s) allowing pre-synaptic C1q expression at the appropriate periods of synaptic pruning in specific brain regions, e.g. the dLGN and the hippocampus (Li et al., 2020; Schafer et al., 2012; Scott-Hewitt et al., 2020).

Augmented release of microglial EVs carrying C1q during active period of synaptic pruning may account for the temporal specificity of C1q association with pre-synaptic terminals. By analysing microglial EVs extracted from the mouse brain across postnatal life, we hereby show that production of Iba-1 and C1q positive EVs peaks at P17, the developmental stage around which synaptic pruning peaks in the hippocampus (Filipello et al., 2018; Li et al., 2020; Paolicelli et al., 2011; Scott-Hewitt et al., 2020; Zhan et al., 2014). This strongly suggests that production of C1q storing EVs by microglia may underly the temporal specificity of C1q association with pre-synaptic terminals during the pruning period in this region. Importantly, we also show that larger production of microglial EVs at peak pruning period in the hippocampus reflects developmental microglial expansion, as evidenced by highest expression of the microglial marker Iba-1 in the mouse brain at P17.

By contrast, production of Iba-1 positive EVs only slightly increases at time of intense pruning in dLGN (P7). However, the dLGN is a small nucleus that accounts for 3.2% of hippocampal volume and about 0.1% of total volume of brain regions used for EVs extraction, according to The Allen Mouse Brain Common Coordinate Framework (Wang et al., 2020). Thus, EVs production from dLGN may marginally influence the amount of EVs that we retrieved. Assessing whether release of microglia-derived EVs is temporally correlated with pre-synaptic C1q localization and pruning in dLGN remains a question for future studies, which requires analysis of EVs uniquely extracted from the nucleus across postnatal development.

### Elevated release of microglial EVs from adult C9orf72 KO brain

Augmented production of EVs from microglia during the pruning period in the postnatal brain raises the question as to whether microglial EVs production represents a general mechanism underlying synaptic tagging by C1q for subsequent microglial removal across the mouse life span.

We hereby show that larger amounts of microglial EVs are produced in C9orf72 KO compared to WT adult brains. This suggests that augmented microglial EVs production and EVs-mediated C1q synaptic deposition may account for excessive microglia-mediated synaptic engulfment in C9orf72 KO brain (Lall et al., 2021). However, we could not prove this hypothesis in the present study, due to the lack of specific tools to control microglial EV release *in vivo.* Therefore, we cannot exclude that other pathological alterations, such as immune system dysregulation, impaired endosomal maturation or lysosomal dysfunction, may contribute to synaptic damage in C9orf72 KO brain at late life stage. Notably, in this context, a strategy to halt hypersecretion of EVs from neurodegenerative microglia has just been published, consisting of silencing of the selenoprotein P (Sepp1)(Bodart-Santos et al., 2024), a regulator of microglial EV biogenesis and release (Ruan et al., 2022). If Sepp1 silencing restores microglial EV production to WT levels in C9orf72 KO mice, this new tool could be used in future studies to clarify whether hyperproduction of EV from C9orf72 KO microglia is causally related to pathological pruning in adult C9orf72 KO mice.

To conclude, our study represents an important step forward in the understanding of how synapses are tagged by C1q for subsequent microglia-mediated synaptic pruning. The identification of microglial EVs as delivery vehicles for C1q to dysfunctional synapses suggests that abnormal EVs production from microglia may be directly implicated in both neurodevelopmental brain disorders and age-related neurodegenerative diseases, in which microglia play a pathogenic role.

## DATA AVAILABILITY STATEMENT

The datasets generated and/or analysed during the current study are available at the following link: https://zenodo.org/records/14937341?preview=1&token=eyJhbGciOiJIUzUxMiJ9.eyJpZCI6ImNjMGEyMjQxLWNjZGEtNGFlZC04YTY5LWUwOTgyNTQ5OGZjNSIsImRhdGEiOnt9LCJyYW5kb20iOiI3N2E3Zjc5MjY2MGMzM2MwM2NjMjE4ZmQxMDE5OThmYSJ9.XD264TY_mJ02fXNp66enDKKSXyuBdsTToBSrAuj_5HKqTj7UmxlubItN-_aoG5WdzQPm_hMqCN-5mjxjJDN0Kg

## ACKNOWLEDGMENTS

We thank Alessandro Gori for precious help and suggestions, Marta Tiffany Lombardo and Camilla Halimi for help with image acquisition and brain EVs extraction, Katia Monsorno for support with Imaris analysis, and Pasquale D’Acunzo for helpful discussion on brain EVs isolation. Cryo-EM was carried out at NOLIMITS, an advanced imaging facility established by the University of Milan.

## FUNDING

This work was supported by RF-2016-02361492 from the Italian Ministry of health to R.G. and C.V., PRIN 2022-3CHCY to C.V., by Regione Lombardia, Italy, POR FESR 2014–2020 resources HUB Ricerca Innovazione project, ID1157625 to C.B., by Next Generation EU through MUR-PRIN 2022 project EV-PRINT (2022CS9H53) to P.G. and IBRO/PERC scientific grant OCT/2022 to G.D..

## AUTHOR CONTRIBUTIONS

Conceptualization C.V., R.C.P.; Methodology G.D., M.G.; Formal Analysis G.D., M.T.G., M.L., M.G.; Investigation G.D., G.C., M.T.G., F.S., M.L., R.F., M.C., P.G., E.B., S.F.C., C.B., S.B., C.S., M.G.; Resources E.A., C.B.; Data Curation G.D., M.T.G., M.G.; Writing – Original Draft C.V.; Writing – Review & Editing G.D., M.T.G., E.A., R.G., C.B., R.C.P., M.G.; Visualization G.D., M.T.G., M.G., C.V.; Supervision G.D., M.C., R.C.P., M.G., C.V.; Funding Acquisition R.G., C.V..

## DECLARATION OF INTERESTS

The authors declare no competing interests.

## METHODS

### Study design

The sample size was estimated based on similar studies previously carried out in the laboratory of others for analysis of synaptic density and microglia engulfment and EVs characterization. Researchers performing the final analysis were blind to the treatment groups. No exclusion criteria were pre-determined. Samples/data points were excluded from the analysis only if they were identified as outliers using a ROUT test (Q = 1%).

### C9orf72 knockout mice (C9orf72 KO)

C9orf72 knockout mice were created by flanking exons 3 and 4 with LoxP sites and crossing them with CMV-Cre deleter mice. The resulting heterozygous offspring were interbred to obtain homozygous knockout (C9orf72 KO) and wild-type controls (WT) on a C57BL/6J background. PCR confirmed exon deletion, and real-time PCR verified the absence of C9orf72 expression, as detailed by Bauer et al. (Bauer et al., 2022). Procedures involving animals and their care were conducted under the institutional guidelines at the Mario Negri institute for pharmacological research IRCCS, Milan. The guidelines comply with national (D.lgs 26/2014; authorization n.493/2019-PR issued on July 4, 2019, by Ministry of Health) and Mario Negri institutional regulations and Policies providing internal authorisation for persons conducting animal experiments (Quality Management System certificate—UNI EN ISO 9001:2008—reg. N° 6121), the NIH Guide for the Care and Use of Laboratory Animals (2011 edition) and EU directives and guidelines (EEC Council Directive 2010/63/UE). All animals were housed under specific pathogen-free (SPF) conditions at a temperature of 22 ± 1 °C, a relative humidity of 55 ± 10% and 12-h light/dark cycle, 5 per cage. Food (standard pellets) and water were supplied ad libitum.

### Cdh5-CreER^T2^::R26^tdTomato^ transgenic mice

Cdh5-CreER^T2^ (Wang et al., 2010) and R26^tdTomato^ (Madisen et al., 2010) transgenic mice were previously described. *Cdh5-CreER^T2^* in C57BL/6/FVB mixed background (F1) female aged 6 to 16 weeks were subjected to overnight timed mating with *R26^tdTomato^* in pure C57BL/6 background. Successful mating was judged by the presence of vaginal plugs the morning after, which was considered 0.5 days post conception (E0.5). A single dose of 37.5 mg/kg of 4-hydroxytamoxifen (4-OHT) dissolved in corn oil was delivered by intra-peritoneal (i.p.) injections to pregnant females at E7.5. To counteract adverse effects of 4-OHT on pregnancies, 4-OHT solutions were supplemented with progesterone (18.75 mg/kg). Mice were housed with free access to food and water at the San Raffaele Scientific Institute and University of Milan Institutional mouse facilities. All experiments were performed in accordance with experimental protocols approved by San Raffaele Scientific Institute and University of Milan Institutional Animal Care and Use Committees (IACUC).

### Resources

Details regarding antibodies, chemicals and supplements are reported in Reagents and Tools Table.

### Primary cultures

All the experimental procedures to establish primary cell cultures followed the guidelines defined by the European legislation (Directive 2010/63/EU), and the Italian Legislation (LD no. 26/2014). Mixed glial cultures were established in accordance with our previous publication (Gabrielli et al., 2022) from 2-day-old (P2) WT and constitutive C9orf72 KO C57Bl/6 mouse pups of either sex, generated and maintained at the Mario Negri Institute of Milano, Italy. Briefly, after dissection, hippocampi and cortices were dissociated by fragmentation with a pipette. Dissociated cells were plated on poly-L-lysine-coated (Sigma-Aldrich, St. Louis, MO, USA) T75 flasks in minimal essential medium (MEM, Invitrogen, Life Technologies, Carlsbad, CA, USA) supplemented with 20% foetal bovine serum (FBS) (Gibco, Life Technologies, Carlsbad, CA, USA), glucose (5.5 g/L, Sigma-Aldrich, St. Louis, MO, USA) and Granulocyte-Macrophage Colony-Stimulating Factor (GM-CSF) from murine GM–CSF–transfected X63 cells (Zal et al., 1994) to promote microglial proliferation. Microglia were harvested from 10-day-old cultures by orbital shaking (200 rpm) and plated on poly-DL-ornithine-coated (Sigma-Aldrich, St. Louis, MO, USA) 60 mm tissue cultured treated petri or 16 mm glass slides and maintained for 48 hours before use. Hippocampal neurons were established from the hippocampi of 18-day-old foetal WT C57Bl/6 mouse embryos of either sex (Charles River, Lecco, Italy) as we described (Gabrielli et al., 2022). Briefly, dissociated cells were plated onto poly-L-lysine treated 24 mm coverslips and maintained in Neurobasal plus with 2% B27 plus supplement (Invitrogen, Carlsbad, CA, USA), antibiotics and glutamate (Sigma-Aldrich, St. Louis, MO, USA).

### EVs isolation and labelling with mCLING-ATTO

We isolated EVs released by primary microglia as we previously described (Prada et al., 2018). 1X10^6^ cells plated on a 60 mm petri were exposed to 1 mM ATP (Sigma-Aldrich, St. Louis, MO, USA) for 30 min in 2ml of Krebs-Ringer’s HEPES solution (KRH) (125 mM NaCl, 5 mM KCl, 1.2 mM MgSO_4_, 1.2 mM KH_2_PO, 2 mM CaCl_2_, 6 mM D-glucose, 25 mM HEPES/NaOH, pH 7.4). Conditioned KRH was collected and pre-cleared from cells and debris by centrifugation at *300× g* for 10 min (twice) (Eppendorf 5810R). Total EVs for Western blotting and Tunable Resistive Pulse Sensing analysis (TRPS) were pelleted at *100000x g* for 1 hour at 4°C (Beckman Optima TL). Large EVs to be supplemented to neurons were instead pelleted by centrifugation at *10000× g* for 30 min at 4°C (VWR CT15FE, rotor Hitachi T15A61), resuspended in 150 µl of conditioned neuronal medium and added to about 1X10^5^ neurons plated on a 24 mm coverslip. In a set of experiments, large EVs were labelled with mCLING-ATTO (Synaptic Systems, Goettingen, Germany), as in (D’Arrigo et al., 2021; Gabrielli et al., 2022). Briefly, large EVs isolated from 1X10^6^ microglia were resuspended in 250 µl of PBS (Phosphate buffer solution) and incubated with 400 nM mCLING-ATTO647 for 5 min in a black tube on ice. The reaction was quenched by adding 250 µl (1:1) of 1% BSA in PBS, dilute with 10 ml of PBS and EVs were re-pelleted at *10000× g* for 30 min at 4°C (Beckman Optima L-90K). Pellet was resuspended in 150 µl of conditioned neuronal medium and spotted on about 1.7X10^5^ neurons plated on a glass coverslip for 3 hours before fixing cells for 6 minutes in 4% paraformaldehyde (PFA) 4% Sucrose. In a set of experiments neurons were pre-treated with Annexin V (8.4 µg/ml) for 15 minutes to cloak externalized PS before incubation with microglial EVs. Fixed neurons were washed in PBS, incubated in Goat Serum Dilution Buffer (GSDB)/Triton X-100 blocking solution for 30 minutes and stained with the following primary antibodies: rabbit anti-C1q (1:200, Abcam, cat.182451), guinea pig anti-Vglut-1 (1:500, Synaptic System, cat.135304) and rabbit anti-Shank2 (1:500, Synaptic System, cat.162202) or mouse anti-PSD95 (Post Synaptic Density protein 95, 1:200, Merck Millipore, MAB1596), diluted in GSDB/Triton and followed by fluorochrome-conjugated secondary antibodies: anti-Rabbit Alexa Fluor 555, anti-guinea pig 488 or anti-mouse 488. Coverslips were mounted on microscope slides and images were acquired with a Zeiss LSM800 confocal microscope equipped with a 63X oil objective. The percentage of puncta double positive for C1q and synaptic markers over total C1q puncta were quantified per field. mCLING EVs-associated C1q puncta colocalizing or juxtaposed to synaptic terminals over total C1q puncta per field were also analysed.

### Immunostaining of microglia

WT and C9orf72 KO microglia were plated on poly-DL-ornithine treated 16 mm glass coverslips at a density of 1X10^5^ cells/coverslip. After 48 hours cells were washed with PBS and live stained with rat anti-CD11b-PE (1:100, BD Pharmigen) or Isolectin IB4-488-conjugated (1:100, Invitrogen, cat. I21414) for 5 minutes before fixation in 4% PFA 4% Sucrose for 15 min. Cells were then washed with PBS, incubated with GSDB/Triton blocking solution for 30 min at RT and stained overnight with primary antibodies: rat anti-Lamp1 (1:100, Invitrogen, cat.14-1072-82), rat anti-CD68 (1:200, Biolegend, clone FA-11), rabbit anti-Cathepsin D (1:400, from (Padovano et al., 2009)), rabbit anti-C1q (1:200, Abcam, cat.182451) diluted in GSDB/Triton followed by species-specific fluorochrome-conjugated secondary antibodies: anti-rat Alexa Fluor 488 or anti-Rabbit Alexa Fluor 555 and DAPI (4′,6-diamidino-2-phenylindole, Invitrogen). Images were acquired with a Zeiss LSM800 with a 40X oil objective and processed with ImageJ software.

### EVs analysis by TRPS

Production of C9orf72 WT and KO EVs was evaluated through Tunable Resistive Pulse Sensing (TRPS) technique, using a qNano instrument (IZON, Christchurch, New Zealand) which provides accurate size and concentration measurements of EVs by detecting current blockage from EVs passing through a nanopore. Recordings have been performed as in (Gabrielli et al., 2022). Cell supernatants from 1X10^6^ microglia WT and C9orf72 KO were cleared from cell debris (*300x g* x 10 minutes twice). Samples were centrifuged at *2000x g* for 20 min, to eliminate bigger EVs, before being pelleted at *100000x g*, resuspended and analysed using an NP200 pore and CPC200 calibration beads. Brain EVs samples were also analysed using NP200 pores and CPC200 calibration beads.

### Western Blotting

Western blotting was performed as in (D’Arrigo et al., 2021). Microglia were lysed with a buffer containing 290 mM sucrose, SDS 1%, 62.5 mM Tris-HCl pH6.8 and proteases inhibitor. A modified version of the Laemmli buffer (15% SDS, 575 mM sucrose, 325 mM Tris-HCl pH6.8, 0.5% β-mercaptoethanol, 0.01% bromo-phenol blue) was then added to a final 1× concentration. EVs from 6-12X10^6^ cells were lysed with the Laemmli buffer. Total lysates of microglia (2.5 µg) and the whole pellet of EVs were loaded on a polyacrylamide gel. Proteins were then separated by electrophoresis, blotted on nitrocellulose membrane and probed for the following antibodies: mouse anti-Lamp1 (1:1000; Invitrogen, cat.14-1071-85), rat anti-CD68 (1:500, Biolegend, cat. 137001 clone FA-11), rabbit anti-Cathepsin D (1:400; Custom), mouse anti-Tmem119 (1:500; Proteintech, cat.66984-1-IG), rabbit anti-C3 (1:500; Abcam, cat.Ab200999), rabbit anti-C1qA (1:500; Invitrogen, cat.PA5-29586), rabbit anti-Alix (1:500; Covalab, cat.PAB0204-0X), mouse anti-Flotillin (1:1000, BD Biosciences, cat.610820), rabbit anti-Annexin A2 (1:5000, Abcam, cat.ab418003), mouse anti-GS28 (1:1000; BD Bioscience, cat. 611184), rabbit anti-GAPDH (1:1000; Synaptic Systems, cat. 347002), rabbit anti-Iba-1 (1:500; Wako, cat. 016-20001), mouse anti-ALDH1L1 (1:500; Millipore, cat. MABN495), mouse anti-GFAP (1:1000; Merck, cat. G3893), mouse anti-L1CAM (1:500; Invitrogen, cat. 14-1719-83), mouse anti-βIII-tubulin (1:2000; Biolegend, cat. MMS-435P), rabbit anti-PLP (1:500, Merck, cat. SAB2101830), rabbit anti-VGLUT1 (1:5000, Synaptic Systems, cat. 135 303), rabbit anti-GM130 (1:1000; kindly provided by Dr M. Renz from the Institute of Immunology and Molecular Genetics, Karlsruhe, Germany (Seelig et al., 1994)) and rabbit anti-Tom-20 (1:500; Santa Cruz Biotechnology, cat.sc-11415). Photographic development was by chemiluminescence (ECL, Euroclone; Westar One Plus, Cyanagen; FEMTO, Thermo Fisher Scientific; or Westar Hypernova, Cyanagen) according to the manufacturer’s instructions. Western blot bands were quantified by ImageJ software.

### Microglia-neuron cocultures

Primary WT or C9orf72 KO microglia were plated on 13DIV hippocampal neurons grown on glass coverslips in a microglia-to-neurons ratio of 1:1. Before microglia coculturing, a set of neurons were treated with EVs isolated from 1X10^6^ 2-day-old WT microglia for 3 hours.

To inhibit EVs release, one T75 cm^2^ flask containing WT or KO mixed astrocyte-microglia was incubated O/N with 5µM GW4869. After treatment, microglia were harvested by shaking, pelleted at *300x g* and plated on neurons. Cells were cocultured for 24 hours at 37°C and 5% CO_2_, fixed in PFA 4% containing Sucrose 4% for 6 minutes, permeabilized in GSDB/Triton blocking solution and immunolabelled for isolectin IB4-488-conjugated (1:100, Invitrogen, cat.I21414), guinea pig anti-VGLUT1 (1:500, Synaptic System, cat.135304), or rabbit anti-Shank2 (1:500, Synaptic System, cat.162202) or guinea pig anti-Bassoon (1:500, Synaptic System, cat.141004) and fluorochrome-conjugated secondary antibodies: anti-rabbit Alexa Fluor 555, anti-guinea pig Alexa Fluor 633 and DAPI. Images were acquired with a Zeiss LSM800 confocal microscope and a 63x oil objective for ImageJ analysis of synaptic density and/or microglia synaptic engulfment.

### Beads Phagocytosis Assay

Fluorescent latex beads (Lian et al., 2016), 1μm in size (Latex beads, Sigma-Aldrich), were pre-opsonized in FBS (Gibco) at a 1:5 beads:FBS ratio at 37°C for one hour, with frequent agitation. Beads were then diluted with MEM (Invitrogen) to reach a final concentration of beads in MEM of 0.01% (v/v) and FBS in MEM of 0.05% (v/v).The medium of microglia at 2DIV, plated on 12 mm slides at a density of 5x10^4^ cells/slide was replaced with 250 μl of bead suspension for one hour at 37°C. Microglial cells were then washed with sterile PBS, fixed with 4% PFA containing Sucrose 4% for 15 minutes washed three times with PBS and stored at 4°C. Cells were labelled with Isolectin IB4-488-conjugated antibody and DAPI. Images were acquired at 40x magnification using a Zeiss LSM800 confocal microscope and analysed with ImageJ.

### Synaptosomes isolation from mouse brain and microglia engulfment

Synaptosomes were prepared as we did in (Tonoli et al., 2022), and according to 10.1038/nprot.2008.171. Cortices were isolated from mouse adult brain and separately disaggregated into a glass potter homogenizer filled up with TS-solution (0.32M sucrose, Tris-HCl 10 mM at pH7.4) in a ratio of 20 ml/g of cortex. After 15 gentle strokes, the homogenate was moved in a 50 ml tube and centrifuged at *1000x g* for 5 minutes at 4°C. Synaptosomes were purified on a three-steps Percoll gradient (23%, 10%, 3%) by centrifuging at *33000x g* for 5 minutes at 4°C (Beckman Optima Max-XP Ultracentrifuge). The band between the 23% and 10% Percoll steps containing synaptosomes was collected and synaptosomes were washed in KRH and pelleted in a tube at *20000x g* at 4°C for 10 minutes. Synaptosomes from 1 cortex were divided in two vials and resuspended in 0.1M Na_2_CO_3_ (about 1mg of synaptosomes in 300µl) and labelled by adding 3 µl of pHrodoTM red dye (previously prepared following manufacturer instructions; Thermo Fisher, Life Technologies). Tubes were mildly vortexed and kept on an orbital shaker at 40 rpm for 2 hours at room temperature, protected from light. After 2 washes with 1 ml of PBS and centrifugation, synaptosomes were resuspended in 300 µl of glial medium, incubated or not with EVs from 1x10^6^ microglia for 3 hours and added to 1X10^5^ microglia plated on 16 mm coverslips. After 2 hours at 37°C and 5% CO_2_, cells were washed 3 times with warm KRH, fixed for 15 minutes in PFA 4% Sucrose 4% and stained with IB4-488-conjugated (1:100, Invitrogen cat. I21414) and DAPI. In a set of experiments, not labelled synaptosomes were exposed to microglia and their engulfment revealed with guinea pig anti-VGLUT1 (1:500, Synaptic System, cat.135304) followed by anti-guinea pig Alexa Fluor 633. Images were acquired at 40X magnification with a Zeiss LSM800 microscope and synaptosome engulfment analysed with ImageJ.

### Immunohistochemistry

P17 male mice were anesthetized and transcardially perfused with 4% PFA in 0.1M PBS and postfixed overnight with 4% PFA in 0.1M PBS. Brains were then kept in a solution containing 30% sucrose and 0.1M PBS until precipitation and then moved to a solution containing cold N-pentane and stored at –80°C till use. Brains were cut in 40 µm thick coronal sections which were stained with the following antibodies: guinea pig anti-Vglut-1 (1:500, Synaptic System, cat.135304), mouse anti-PSD95 (Post Synaptic Density protein 95, 1:200, Merck Millipore, MAB1596), rabbit anti-C1q (1:200, Abcam, cat.182451), rabbit anti-Iba1 (1:200, Wako, cat.019-19741) and rat anti-CD68 (1:200, Biolegend, cat. ab53444). Primary antibodies were incubated overnight at 4°C in 10% normal goat serum with 0.3% Triton-X 100 in PBS. Sections were then exposed for 2 hours at room temperature to species-specific secondary antibodies: anti-guinea pig Alexa Fluor 488, anti-mouse Alexa Fluor 488, anti-rabbit Alexa Fluor 555, anti-rabbit Alexa Fluor 633 and anti-rat Alexa Fluor 555. Finally, after a 5-minute incubation with DAPI (1:10000), sections were mounted with DAKO fluorescent mounting medium (Agilent, cat.S3023). Images were acquired with a 63X oil objective mounted on a Zeiss LSM800 confocal microscope with a z-step size of 0.33µm and analysed with ImageJ or Imaris.

### OT manipulation

Hippocampal neurons at 10-12 DIV plated on glass coverslip were transfected with 0.35 µg of farnesyl-GFP (f-GFP) and Vglut1-mCherry expressing vector using Lipofectamine 2000 to visualize neuron morphology and delineate dendritic spines receiving VGLUT1 pre-synaptic puncta. After 48-72 hours, neurons were washed and mounted in 500 µl of KRH on a temperature controlled live imaging chamber of an inverted spinning-disk confocal microscope (Axiovert 200 M, Zeiss, Oberkochen, Germany), equipped with an IR laser beam (1064 nm, CW) collimated into the optical path of microscope lens (Zeiss 63X objective, oil immersion, NA 1.4). Large EVs isolated by centrifugation at *10000x g* from about 1X10^6^ microglia were resuspended in 250µl of KRH and 25µl of the EVs suspension was added to about 1X10^5^ neurons. Single EVs which appeared in the recording field were trapped and positioned on selected f-GFP+ dendrites nearby VGLUT1 spots, as previously described^22^. After adhesion, EVs were monitored by time lapse imaging for 30 minutes. The initial and final EVs positions on processes were imaged and the number of EVs that localized at VGLUT1 puncta (colocalizing or juxtaposed) was quantified.

### Image analysis in microglia *in vitro*

The signal intensities of CD68, Lamp1, Cathepsin D and CD11b were quantified with ImageJ by using IB4 to highlight individual cell area. After setting the optimal ‘Color Balance’ values for the channels of interest (same values for all conditions) and collecting the cell area, the integrated intensity and the mean grey values of CD68, Lamp1, Cathepsin D and CD11b, the following formula was used to calculate the corrected total cell fluorescence (CTCF) = integrated density – (area of selected cell × mean fluorescence of background readings) (Zhang et al., 2020).

### Analysis of pre- and post-synaptic puncta in microglia-neuron cocultures

Pre- and post-synaptic density of neurons cocultured or not with microglia was analysed with ImageJ software. Optimal ‘Color Balance’ and ‘Threshold’ values were defined for Shank2 and Bassoon channels and applied to all images in the same experiment. Segments of neurites close to microglia with visible synaptic puncta were measured using the ImageJ ‘Segmented Line’ tool. For each segment the number of pre-synaptic and post-synaptic puncta were counted using ‘Multi-point’ function. To calculate the density of puncta, the number of puncta in each channel was divided by the length of the relative segment. Data were normalized on the mean of controls.

### Analysis of microglia engulfment *in vitro*

Maximum projection of Z-stacks was used to quantify synaptic engulfment *in vitro*. IB4 signal was used to define microglial cell area. Optimal thresholds values were defined for synaptic puncta labelled by Bassoon, Vglut-1 or Shank2 or pHrodo-labelled synaptosomes and applied to all experimental conditions. Analysis of engulfed material inside microglia was carried out using the ‘Colocalization’ plug-in in ImageJ. Colocalizing area (IB4 positive synaptic area) per image was collected and normalized on microglia area.

To quantify beads phagocytosis, the number of microglia with engulfed beads over the total number of microglia per field was calculated.

### Analysis of hippocampal slices

Maximum projection of Z-stacks was used for the analysis of C1q, Vglut-1 or PSD95 positive area per field. The same threshold was set for all conditions and total immunoreactive area per field was measured using the ImageJ ‘Analyzed particles’ function. For analysis of C1q colocalization with pre- or post-synaptic puncta, three consecutive stacks per field, with highest C1q signal, were selected. The colocalizing area was calculated in the three stacks using the ImageJ ‘Colocalization’ function.

Analysis of synaptic engulfment in individual microglial cells was carried out by using Imaris software (version 9.9.1, Bitplane). Entire individual microglial cells were 3D reconstructed with the built-in “Surface” function and Iba-1+ area and volumes were quantified using the same thresholds across conditions. Synaptic volumes (Vglut-1 and PSD95) were reconstructed using the Imaris “spot” plugin inside Iba-1 or CD68 volumes, quantified and normalized on total volume of individual Iba-1+ cells.

### ELISA

C1q ELISA Kit (#NBP2-74988, Novus Biologicals) has been employed to measure C1q in EVs and EVs-depleted cell supernatant in accordance with manufacturer instructions, as in (Lombardi et al., 2019). EVs were isolated from 16x10^6^ microglial cells exposed to 1 mM ATP for 30 min in KRH. Cell supernatant containing EVs was supplemented with cOmplete, EDTA-free Protease Inhibitor (#04693132001, Roche) and cleared through two *300x g* 10 min centrifugations. Then, total EVs were pelleted at *100000x g* and resuspended in 100 µl ELISA buffer in the presence of protease inhibitors (1:100, Halt™ Protease and Phosphatase Inhibitor Cocktail, #78440, Life Technologies). The supernatant was then depleted of residual particles by ultracentrifugation at *200000x g* for 3h. Trichloroacetic acid 1:10 v:v was added to the sample, which was inverted several times and left for 1h on ice for protein precipitation before pelleting at *15000x g* for 5 min at 4°C. The obtained pellet was washed with ice-cold 100% Acetone, re-pelleted and resuspended in ELISA buffer with protease inhibitors (Halt). Just before loading in the ELISA plate, 0.6% Triton X-100 (0.6 µl, Sigma) were added to EVs samples. The same amount of Triton X-100 and protease inhibitors was added to the supernatant sample, and to standard dilutions used to estimate C1q amount based on a calibration curve.

### Cryo-EM

Freshly prepared EVs released from 10 million microglia, and frozen EVs from adult brain were resuspended in saline/PBS and vitrified by applying a 3.0 μl droplet onto holey carbon grids (Copper 300-mesh Quantifoil R2/1) or Lacey carbon Films (+2nm 400-mesh), glow discharged for 45 seconds at 40 mA, or for 30 seconds at 30 mA, using a GloQube system (Quorum Technologies, East Sussex, UK). After 60/90 seconds incubation, the grid was plunge-frozen in liquid ethane using a Vitrobot Mk IV (Thermo Fischer Scientific, Waltham, MA, USA) operating at 4◦C and 100% RH. Images of the vitrified specimen were acquired using a Talos Arctica transmission electron microscope (Thermo Fisher Scientific, Waltham, MA, USA) operating at 200 kV and equipped with a Falcon 3EC (or a Ceta 16M) direct electron detector (Thermo Fischer Scientific, Waltham, MA, USA). Images were acquired with an exposure time of 1 second and a total accumulated dose of 80 electrons per A2 at a nominal magnification of 45000 × or 36000 x, with applied defocus values between 2 and 4 μm. Images acquired with Volta phase-plate were recorded at defocus values between 0.5 and 1 μm. Contrast transfer function (CTF) estimation was performed using CTFFIND4 (Rohou and Grigorieff, 2015). This analysis has been carried out at NOLIMITS facility (University of Milan).

### Isolation of EVs from the interstitial fluid of mouse brains

Brain EVs were isolated from mouse brain tissue using a methodology (Fig. 5A) adapted from published protocols (Crescitelli et al., 2021; D’Acunzo et al., 2022; Vella et al., 2017). Briefly, brains were collected from female WT or C9Orf72 KO mice, or Cdh5-CreER^T2^::R26^tdTomato^ transgenic mice of different age after sacrifice in accordance with the European directive (2010/63/EU) and the Italian Law for Care and Use of Experimental Animals (DL116/92; DL26/2014). Brain extraction was performed on dry ice on a glass surface. Tissues were snap frozen in liquid nitrogen and stored at -80°C pending use. To isolate EVs, frozen brains were placed in Hibernate-A (Life Technologies) and meninges rapidly but carefully peeled off. Then, brains were weighted and approximately 320-380 mg of tissue were cut into small pieces (1-2 mm^3^) in 1 ml Hibernate-A on ice, added to a 75 U/ml Collagenase type I (#LS004194, Worthington Biochemical Corporation) Hibernate-A solution and incubated at 37°C for 15 min. After incubation, samples were placed on ice, added with protease and phosphatase inhibitors (#04 693 116 001, cOmplete Protease Inhibitor Cocktail, and #04906845001, PhosSTOP Phosphatase Inhibitor Cocktail, Roche), and filtered using a 70 μm strainer to eliminate not dissociated tissue (MACS SmartStrainers, #130-110-916, Miltenji). The samples were then subjected to differential centrifugation at *300x g* for 10 min and *2000x g* for 20 min at 4°C to clear them from cells and cellular debris. The supernatants were transferred to new tubes and centrifuged at *10000x g* for 30 min at 4°C (rotor Fiberlight F15-8, Sorvall Legend XFR, Thermo Fisher Scientific) to obtain low-speed pellets enriched in high density/large size EVs. The pellets were resuspended in cold PBS, filtered through a 0.45 µm filter (SFCA, #431220, Corning) and re-pelleted at *10000x g* for 30 min at 4°C. In parallel, the *10000x g* supernatants were filtered through a 0.20 µm filter (SFCA, #431219, Corning) and ultracentrifuged at *120000x g* for 130 min at 4°C (26500rpm with rotor Sw 41 Ti, Beckman Optima L-90K), washed and re-pelleted to obtain a high-speed pellet enriched in lower density/smaller size EVs. Crude low-speed EVs (l-EVs) and high-speed EVs (h-EVs) were resuspended either in cold PBS to be stored at -80°C or in 25 mM trehalose/PBS to be preserved overnight before gradient purification. In the latter case, crude l-EVs and h-EVs were carefully loaded on top of iodixanol step density gradients, composed of 4 ml 40%, 3 ml 20%, and 3 ml 5% Optiprep (#D1556, Merck) in 25 mM trehalose/PBS loaded in a bottom-up manner, and ultracentrifuged at 234000x g for 5h (37000 rpm with Sw41Ti rotor, Beckman Optima L-90K). After centrifugation, 5 fractions (2.2 ml each) of density medium were carefully pipetted from top to bottom (1 to 5). Each fraction was transferred into a new tube, diluted in cold PBS and ultracentrifuged at *120000x* g x 130 mins at 4°C. Obtained pellets were resuspended in cold 25 mM trehalose/PBS and stored at -80°C. For Western Blot analysis, EVs samples were digested in modified Laemmli Buffer.

### Flow cytometry analysis

Single cell suspensions were obtained from 3-months old mice telencephalon by incubating for 15 min at 37°C in calcium/magnesium free PBS supplemented with FBS 10%, Penicillin-Streptomycin 1%, EDTA 2mM and collagenase type I (Sigma) 0.12% and dispase 0.15% (w/v). Then samples were treated with 10% (w/v) DNAse in calcium/magnesium free PBS solution for 15 min at 37 °C, followed by mechanical dissociation by pipetting. Single cell suspensions were incubated with mouse anti-CD45 FITC and rat anti-F4/80 PE-CY5 antibodies (Biolegend), rat anti-Tmem119 PE-Cy7 and rat anti-CD31 APC (eBioscience) and processed for flow cytometry as described (Azzoni et al., 2021; Azzoni et al., 2018). Voltages, compensation and gates were set using unstained, single stained and fluorescence-minus-one (FMO) controls. Dead cells were excluded based on Hoechst 33258 (Hellobio). Flow cytometry data acquisition was carried out using a LSR Fortessa X-20 (BD) analyzer and BD FACSDiva software (version 8.0.2). Flow cytometric data was analyzed using FlowJo software version 10 (BD).

### Staining of brain EVs

L-EVs from Cdh5-CreER^T2^::R26^tdTomato^ adult transgenic brain tissue (50 mg) were resuspended in 200 µl of PBS and incubated with rat anti-mouse CD11b-CY5 (1:100, Biolegend, Cat.101209) for 10 minutes. L-EVs were re-pelleted at *10000× g* for 30 minutes at 4°C and resuspended in 200 µl GSDB containing Alexa anti-Rat 647 secondary antibodies (1:200) for 2 hours at room temperature. After further washing in PBS and re-centrifugation as above, the EVs were resuspended in 100 µl of PBS, spotted on a glass coverslip and imaged with a Zeiss LSM800 confocal microscope equipped with a 63X oil objective.

L-EVs from P17 WT brain tissue were incubated for 20 minutes with Calcein-AM (1:750, Life Technologies, Cat.F10322) before adding CD11b-CY5 or O4-CY5 or IB4-568 (1:100, Thermo Fisher, cat. 121412) for further 10 minutes. L-EVs were then re-pelleted as above, resuspended in 200 µl GSDB in the presence of rabbit anti-C1qA primary antibody (1:200, Abcam, cat.182451) or rat anti-C3 (1:100, Abcam, cat. ab11862) and rabbit anti-GLAST C-term (1:100) for 1 hour at room temperature on a wheel, further washed with PBS and re-pelleted before incubation with Alexa anti-rabbit 555 and anti-Rat 647 (1:200) or Alexa anti-rat 555, anti-rabbit 405 and anti-mouse 633 (1:200) for 2 hours at room temperature. After final wash and re-centrifugation, EVs were resuspended in PBS and spotted on glass coverslip for image acquisition. The percentage of puncta double positive for C1q/C3 and CD11b or IB4 or GLAST or O4 over total puncta were quantified.

### Brain homogenates

Brains collected from WT/C9orf72 KO C57Bl/6 mice of different age were homogenized in Lysis Buffer (290mM sucrose, 1% SDS, 0.125% v/v Tris-HCl 0.5M pH6.8; 1ml/100mg) supplemented with protease inhibitor cocktail (#P8340, Merck) using a Teflon-glass Potter. Homogenates were centrifuged at *1600x g* for 15 min, supernatant collected and stored at -80°C. For Western Blot analysis samples were digested in modified Laemmli Buffer.

### SiMoA

SiMoA carboxylated paramagnetic beads conjugated with the bradykinin-derived membrane sensing peptide (MSP), a peptide sequence with affinity for EVs, were used as EVs capture probe to perform a single vesicle analysis on SiMoA platform (Gori et al., 2024). The following three-step assay was performed. The MSP-beads solution was prepared at a concentration of 2x10^7^ beads/mL in Bead Diluent (Quanterix), according to manufacturer’s instructions. EVs were diluted 1:40 in Homebrew Sample Diluent (Quanterix), and 100 µL of the sample was mixed with 25 µL of functionalized beads in a 96-well plate and incubated for 90 minutes at 25°C, under shaking at 800 rpm. After a washing step performed by an automatic plate washer, 100 µL of biotinylated detector antibody (anti-CD11b; eBioscience, cat. 14-0112-82; anti-C1q, abcam, ab182451) was added and incubated for 10 minutes at 25° C, under shaking at 800 rpm. Antibodies were incubated at 0.6 mg/mL in Detector Diluent (Quanterix). The beads were then washed, incubated for 10 minutes with a 150 pM SBG solution (in SBG Diluent, Quanterix), washed again and inserted into the Quanterix SR-X instrument for analysis, where RGP reagent was automatically added. In a set of experiments SiMoA carboxylated paramagnetic beads were conjugated with anti-Cd11b according to manufacturer’s instructions, to capture EVs produced by brain macrophages. Three-step SiMoA assay was performed as described, with the first incubation step lasting for 30 minutes at 25°C, under shaking at 800 rpm, as recommended by manufacturer’s protocols. Data were analysed and processed using SiMoa 1.1.0 Reader Software.

### Statistical analysis

Statistical analysis was performed on all datasets. Data were first tested for normal distribution with GraphPad Prism 6 software, then the appropriate statistical test has been used. The accepted level of significance was p ≤ 0.05, indicated by one asterisk; those at p ≤ 0.01 are indicated by double asterisks, the ones at p ≤ 0.001 are indicated by triple asterisks.

## APPENDIX SUPPLEMENTAL FIGURES

Appendix supplemental figures S1-S10.

